# Cold-responsive MtCBF4-MtJMJ13 positive feedback loop negatively regulates anthocyanin biosynthesis in *Medicago truncatula*

**DOI:** 10.1101/2025.10.18.683229

**Authors:** Niyaz Ahmed, Jogindra Naik, Debasis Chattopadhyay, Ashutosh Pandey

## Abstract

Legumes, being significant food sources for both humans and animals globally, experience several adverse conditions that considerably affect their crop yield. *Medicago sp.*, an important forage crop, thrives under adverse conditions, such as drought, extreme temperatures (heat and cold), and salinity. Several studies have shed light on the regulatory pathways of cold/chilling stress tolerance in *Medicago sp.* in recent years. In the present study, we found that a cold-repressed transcription factor, MtCBF4, negatively regulates anthocyanin synthesis by directly repressing the expression of *MtLAP1*, a critical regulatory gene for anthocyanin synthesis in *Medicago truncatula*. It leads to the downregulation of anthocyanin biosynthesis genes, and reduced anthocyanin accumulation. Apart from the transcriptional cascade, MtCBF4 exerts its effect through histone H3K27 trimethylation. It directly binds to the promoter of *MtJMJ13*, an H3K27me3 demethylase, to activate its expression, resulting in decreased H3K27 trimethylation. Furthermore, MtJMJ13 demethylates the gene body of *MtCBF4* to positively regulate its expression, forming a positive feedback loop that functions in a cold-responsive manner. MtJMJ13 also demethylates the gene body of *MtMYB2*, *MtKFB1*, and *MtKFB2* genes, which are negative regulators of anthocyanin biosynthesis. Altogether, we have unveiled a new regulatory loop involving a transcriptional regulator and a histone demethylase, operating together to limit anthocyanin production. This regulatory module is repressed under cold stress, leading to increased anthocyanin production and enhanced cold stress tolerance in *M. truncatula*.

## Introduction

Abiotic stresses pose a huge threat to the productivity of plants with changing global climates (Chaudhry et al., 2022; Zandalinas et al., 2022). The ability to withstand moderate to severe abiotic stress as a key feature evolved quite early in plants and is mediated by a complex hierarchical regulatory nexus (Zuo et al., 2023; Gill et al., 2024). The plant’s resilience to different abiotic stresses is largely mediated by complex regulatory cascades operating at various levels in sequential manner (Prusty et al., 2024; Saleem et al., 2025). Legume crops, contributing one-third of the total yields, are a source of nutrition for both humans and animals. *M. sativa*, among roughly 87 species of the Medicago genus, is labelled as “queen of forage” and has been extensively grown as a forage crop worldwide (Ye et al., 2025). It plays a vital role in animal husbandry, where it provides a high level of proteins, vitamins, minerals, and a balanced array of amino acids to the livestock (Ye et al., 2025).

Among the major abiotic constraints encountered by plants, drought, high salinity, and cold majorly affect the physiology and productivity (Singh et al., 2018; Zhang et al., 2022). Out of these, cold stress is experienced by the plants when exposed to low temperatures (0- 15L), while temperatures below freezing points (0L) cause freezing stress, which is more deleterious to the plants (Bracale et al., 2003). Legumes like *Medicago sp.* sustain unfavourable low temperature conditions, which impact their productivity, photosynthesis, reproductive development, nodule symbiosis, and ultimately forage yield (Wang et al., 2021; Zhang et al., 2022). To withstand adverse low-temperature conditions, plants become triggered to initiate internal signalling cascade involving several layers of regulatory components. The induction of stress-responsive COR genes (cold shock proteins, transcription regulators, and transporters), as well as the expression of ROS scavenging antioxidant enzymes and flavonoid pigments are major biochemical processes adopted by plants during cold stress (Kidokoro et al., 2022; Gusain et al., 2023). Like other plants, *Medicago sp.* also possesses regulatory pathways that confer cold tolerance, many of which have been revealed in recent studies (Zhou et al., 2018; Adikari et al., 2022). Various aspects of anthocyanin biosynthesis have been studied in recent years; However, how the anthocyanin biosynthesis is modulated under cold stress conditions has not been examined in detail in *Medicago sp*.

Flavonoids group of specialized metabolites encompasses a wide array of different sub-classes like anthocyanin, flavonol, flavone, proanthocyanidin, flavanone, aurone, isoflavone, etc., which vary greatly in their structures and chemical natures (Tohge et al., 2017; Shen et al., 2022; Naik et al., 2022). They play critical roles in the physiology and stress tolerance mechanisms in plants and also impart various health-beneficial functions to the animals by exerting anti-cancer, anti-fungal, antioxidant, and anti-inflammatory properties; thus, these are essential for animal and human health (Taylor et al., 2005; Buer et al., 2010; Agati et al., 2012; Jucá et al., 2020; Yang et al., 2023; Naik et al., 2024). Anthocyanins, among these, are known to be highly induced under abiotic stresses, particularly salinity, drought, and cold stress, as a strong antioxidant molecule (Kovinich et al., 2014; Naing et al., 2021; Buitrago et al., 2025). They exclusively confer vivid coloration to different floral organs of the plants, and thus, are critical for pollination (Naik et al., 2022; Anum et al., 2024). They are mainly found in the form of glycosides like cyanidin, delphinidin, malvidin, pelargonidin, peonidin, and petunidin (Schnitker et al., 2024; Zong et al., 2025). Anthocyanin is synthesized by the late flavonoid biosynthetic pathway genes, deriving flux from the general phenylpropanoid pathway. Several enzymes like chalcone synthase (CHS), dihydroflavonol 4-reductase (DFR), anthocyanidin synthase (ANS), UDP-glucose:flavonoid-O-glycosyltransferase (UFGT), O-methyltransferases (OMTs) catalyze critical steps in the biosynthesis of anthocyanin in plants (Jaakola., 2013; Zhang et al., 2014; Pandey et al., 2016; Naik et al., 2022). The biosynthetic genes and the regulatory mechanisms have been widely studied in different plant species, including early land plants (Albert et al., 2018; Krishnamoorthi et al., 2024; Li et al., 2024; Liu et al., 2025). Many studies have unfolded transcriptional, epigenetic, post-translational, and post-transcriptional regulations governing anthocyanin synthesis (LaFountain et al., 2021; Naing et al., 2021; Li et al., 2023; Bulgakov., 2024). These regulatory factors synergistically modulate anthocyanin accumulation either *via* integrating environmental stimuli or the hormonal signal transduction pathway. Among the regulators, MYB-bHLH-WD40 triads are the most comprehensively studied master transcriptional regulators of anthocyanin synthesis in diverse land plants (Xu et al., 2015; Davies et al., 2020; Naik et al., 2022; Sharma et al., 2024).

CBF/DREB family of transcription factors are critical regulators of cold stress signalling in plants (Zhao et al., 2016; Shi et al., 2018; Song et al., 2021). They target specific core RCCGAC motifs, also called DRE/CRT (Dehydration Responsive Element/C-repeat), in their target gene promoters (Sukuma et al., 2002; Gierczik et al., 2017; Nie et al., 2022). Many studies have earlier reported diverse roles of these regulatory factors in cold signalling, where these are known to modulate ROS homeostasis, antioxidant metabolism, circadian rhythm, flowering, metabolite synthesis, etc (Shi et al., 2016; Han et al., 2020; Cui et al., 2025). As a terminal regulator in cold signaling, CBF/DREBs are also reported to regulate anthocyanin and flavonoid biosynthesis. For example, BcCBF2 directly binds to the DRE in the promoter of *BcMYB111* to induce its expression in *Brassica sp*. BcMYB111 further regulates the expression of *BcF3H* and *BcFLS1* to modulate flavonoid biosynthesis in a cold stress-dependent manner (Chen et al., 2023). In eggplant, SmCBF1/2/3 interact with SmMYB113 via their C-terminal to promote the transcription of *SmCHS* and *SmDFR*, leading to increased anthocyanin accumulation under cold stress (Zhou et al., 2020). Apart from these, CBF/DREB and anthocyanin synthesis are also coregulated at the transcriptional level by upstream transcription factors of cold signalling (Xu et al., 2023; Zhu et al., 2025). In Medicago sp., few CBF/DREB genes have been demonstrated to play roles in growth, abiotic stress tolerance, flowering, and lignin metabolism (Li et al., 2011; Zhang et al., 2026; Cui et al., 2025).

In the epigenetic regulatory mechanism, histone modification serves as the key player in activation or repression of gene expression through orchestration of interchangeable euchromatin-heterochromatin status (Fuchs et al., 2006; Liu et al., 2022). Histone methylation is the addition of a methyl group onto the lysine/arginine residue of histone proteins by *S*-adenosyl-methionine-dependent methyltransferases (lysine methyltransferases, KMTs, and protein arginine methyltransferases, PRMTs), whereas demethylation is carried out by histone lysine demethylase (KMDs), either flavin-dependent KMDs or Fe(II)- and 2- oxoglutarate (2OG)-dependent JmjC-domain-containing proteins (Liu et al., 2010; Xiao et al., 2016; Hu and Du., 2022). This writer and eraser components tend to maintain histone lysine methylation in the nucleus (Xiao et al., 2016; Hu and Du., 2022). Different methylation contexts of histones, particularly of H3, result in either activation or deactivation. For example, H3K4me3 and H3K36me2/3 are associated with transcriptionally active state, and H3K9me2/3 and H3K27me3 are associated with repressive state (Liu et al., 2010; Xiao et al., 2016). Histone demethylases are classified broadly into five groups: ARID1/KDM5, JHDM3/KDM4, JHDM2/KDM3, JHDM6, and a class of proteins only containing the JmjC domain (Lu et al., 2008). Many histone writers and erasers have been characterized in Arabidopsis and a few crop plants regulating different processes like abiotic stresses (drought, cold, salinity, heavy metal, nutrient starvation), flowering, thermomorphogenesis, flavonoids synthesis, fruit ripening etc. (Fan et al., 2018; Huang et al., 2019; Li et al., Li et al., 2022; Zhao et al., 2025; Yu et al., 2025). Jumonji -domain containing proteins have been largely studied as epigenetic regulators so far, regulating diverse kinds of physiological processes in Arabidopsis and a few more plant species (Dutta et al., 2017; Huang et al., 2019; Zheng et al., 2019; Li et al., 2020; He et al., 2022; Zhao et al., 2025). Arabidopsis genome encodes 21 JmjC domain-containing genes, which are known to control DNA methylation, hormone signaling pathway, bud regeneration, RNA silencing, etc. (Lu et al., 2008; Yu et al., 2008; Audonnet et al., 2017; Fan et al., 2024). *M. truncatula* genome encodes 34 putative JMJ domain-containing proteins, which have been further divided into 5 sub-families (Lopez et al., 2022).

*M. sativa*, an autotetraploid (2n = 4x = 32), possesses many complexities like a large genome size (800-1000 Mb), perennial growth, and high rates of heterozygosity, making its close diploid relative *M. truncatula* a convenient model plant for biological research (Ye et al., 2025). The latter has a small genome (2*n* = 16, ∼430 Mb) and possesses certain traits like self-fertilization, annual growth, a short life cycle, and high genetic transformation efficiency etc. Many studies on growth, development, and abiotic stress resilience have surfaced in recent years, which can be a good toolkit for breeding in *Medicago sp.* In our current study, we have discovered a novel positive feedback regulatory loop involving MtCBF4, a transcription regulator, and MtJMJ13, a histone demethylase, which acts cooperatively to limit anthocyanin production in *M. truncatula* under normal growth conditions, while cold stress represses expression of both genes to enhance anthocyanin production in *M. truncatula*.

## Results

### MtCBF4 negatively regulates anthocyanin accumulation in *M. truncatula*

MtCBF4 is known to confer salt and freezing tolerance in *M. truncatula* and drought stress tolerance in Arabidopsis (Li et al., 2011; Zhang et al., 2016). We observed that two putative chalcone synthase (CHS) genes and a putative Kelch repeat F-box gene, which is known to degrade CHS enzyme in Arabidopsis, are differentially expressed in the comparative transcriptome data of wildtype (WT) and *MtCBF4* overexpressing transgenic lines of *M. truncatula* (Zhang et al., 2016). To examine the role of MtCBF4 in anthocyanin biosynthesis, we transiently overexpressed *MtCBF4* in *M. truncatula* A17 accessions. The expression of the *GFP* in the transformed leaves, as determined through semi-quantitative PCR, confirms successful transformation of the overexpression cassette (Fig. S1). First, we analysed the expression of *MtCBF4*, and observed ∼8-fold increase in its transcript level, suggesting moderate overexpression (Fig. 1A). Further, we analysed the expression of flavonoid biosynthetic genes, which revealed a significant decrease in the expression of many early and late biosynthesis genes, i.e., *MtPAL, MtCHS, MtCHI, MtFLS1, MtF3’H, MtDFR1, MtANS, MtANR, MtLAR, MtGSTF7* etc (Fig. 1A, Fig. S2). *MtGSTF7*, a homolog of Arabidopsis TT19, is crucial for the accumulation of anthocyanin (Wang et al., 2022). *MtGSTF7* expression was reduced drastically in the *MtCBF4* overexpressing leaves. This intrigued us to check the expression of the regulatory genes, and as expected, the expression of *MtLAP1*, *MtTT8* decreased drastically in the *MtCBF4*-overexpressing leaves, while no changes were observed in the case of *MtWD40-1 (Fig. 1A, Fig. S2)*. These three genes are the representatives of MYB-bHLH-WD40 complex members known to regulate anthocyanin synthesis in *M. truncatula*. One putative Kelch repeat F-box gene, *Medtr5g033880*, was differentially expressed in *MtCBF4* overexpressing tissues (Zhang et al., 2016). It is a putative ortholog of KFB^CHS^ known to degrade CHS in Arabidopsis (Zhang et al., 2017). We did phylogenetic analysis and found that the *M. truncatula* A17 genome encodes three putative orthologs of KFB^CHS^, which form a cluster together with AtKFB^CHS^ (At1g23390) (Fig. S3). The three genes encoded by the locus *Medtr8g069800*, *Medtr5g043750*, and *Medtr5g033880* were named as *MtKFB1*, *MtKFB2*, and *MtKFB3*, respectively, in the order of their sequence identity to their Arabidopsis homolog. Expression analysis of these genes in *MtCBF4*-overexpressing tissues showed a significant increase in the expression of *MtKFB1* (3.5-fold), followed by *MtKFB2* (∼3-fold), and *MtKFB3* (∼2-fold) (Fig. 1A). *MtMYB2*, a homolog of *AtMYBL2*, is a known repressor of anthocyanin biosynthesis in *M. truncatula* (Jun et al., 2015). We noticed that *MtMYB2* is highly up-regulated in *MtCBF4-*overexpressing leaves (Fig. 1A). These results suggest a negative regulatory role of MtCBF4 in anthocyanin biosynthesis. Finally, we quantified the anthocyanidin content in *MtCBF4-*overexpressing tissues, which was found to be decreased significantly (Fig. 1B). LC-MS-based metabolomic profiling further verified the decrease in the anthocyanin levels, particularly cyanidin, malvidin, pelargonidin, peonidin, and petunidin (Fig. 1B).

**Figure 1.**
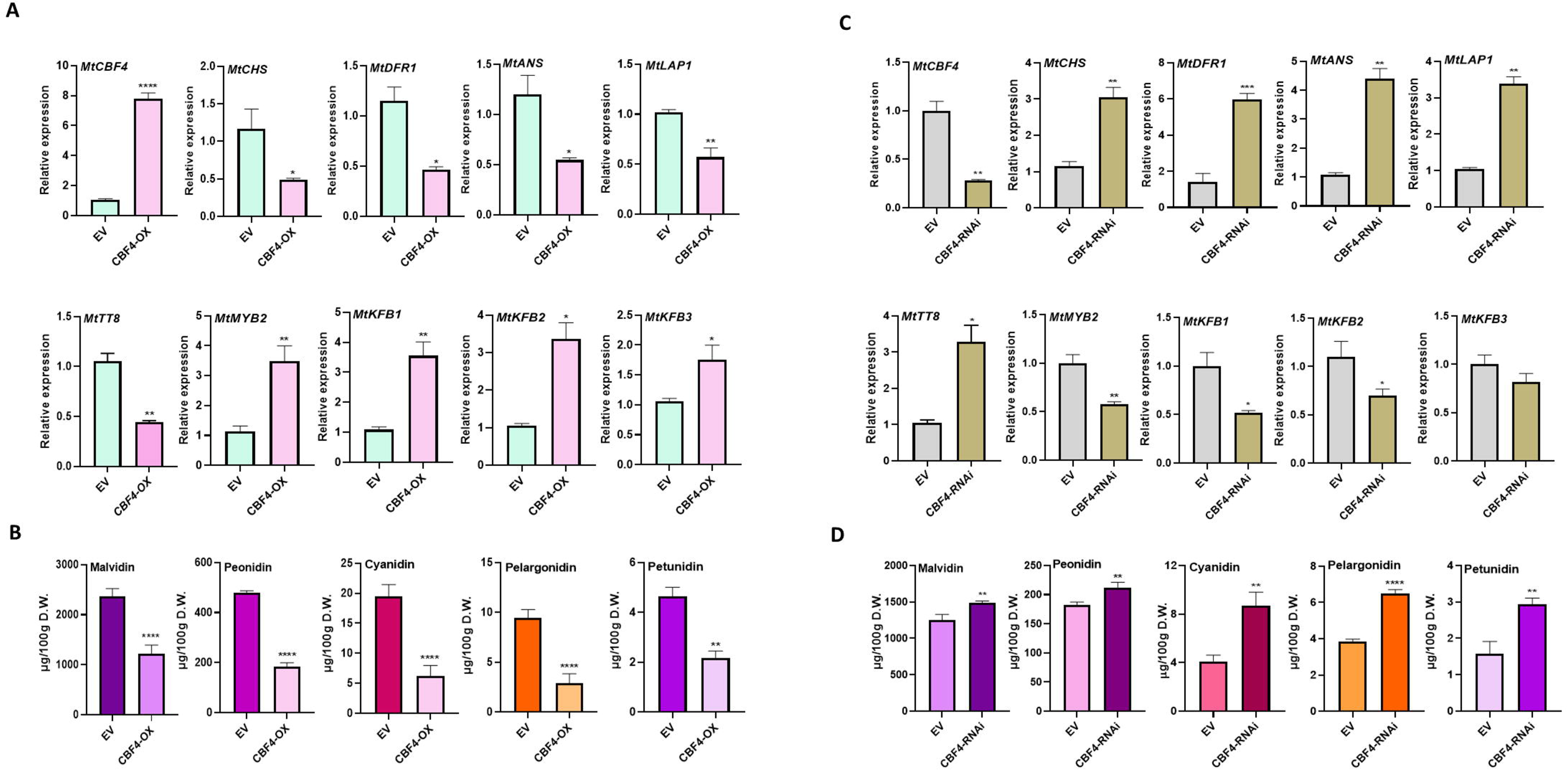
*MtCBF4* represses anthocyanin accumulation in *M. truncatula*. **(A)** Relative expression levels of *MtCBF4*, biosynthesis genes (*MtCHS*, *MtDFR1*, *MtANS*), regulatory genes (*MtLAP1*, *MtTT8*, *MtMYB2*, *MtKFB1/2/3*) in leaves transiently overexpressing *MtCBF4*, as determined by RT– qPCR. *MtACTIN* was used as the internal control. **(B)** Quantitative targeted LC-MS-based quantification of major anthocyanidins (cyanidin, malvidin, peonidin, pelargonidin and petunidin) content in leaves that transiently overexpress *MtCBF4*. The data are mean values ±SD (n=3 biological), each derived from leaves of several plants. **(C)** Relative expression level of *MtCBF4*, biosynthesis genes, and regulatory genes in transformed hairy roots silencing *MtCBF4*. **(D)** Targeted LC–MS analysis of major anthocyanidins content in transformed hairy roots silencing *MtCBF4*. The data are mean values ±SD (n=3 biological), each derived from several hairy roots.

To further corroborate our findings, we generated hairy root RNAi lines of *MtCBF4* in *M. truncatula* A17 ecotype. Bright red fluorescence of transformed hairy roots under a fluorescence microscope due to the expression of the marker gene DsRed confirmed successful transformation of the gene silencing cassette (Fig. S4). Expectedly, the expression of *MtCBF4* significantly decreased in the *MtCBF4*-silenced hairy roots (Fig. 1C). Many biosynthetic genes were significantly up-regulated along with the regulatory genes like *MtLAP1,* and *MtTT8* (Fig. 1C, Fig. S5). Whereas *MtKFB1, MtKFB3,* and *MtMYB2*, which are the probable and established negative regulators of anthocyanin biosynthesis, respectively, were down-regulated in the *MtCBF4*-silenced hairy root tissues (Fig. 1C). Further LC-MS-based targeted metabolite analysis showed an increase in the content of all the major anthocyanidins (Fig. 1D). Altogether, these results establish the strong negative regulatory role of MtCBF4 on anthocyanin synthesis in *M. truncatula*.

### MtCBF4 directly binds to the promoter of *MtLAP1* and represses its expression

To investigate the direct regulatory role of MtCBF4 on pathway genes, we conducted a promoter analysis of key biosynthesis genes that showed significant modulation in the expression level. We searched for the DRE/CRT element (RCCGAC), the binding motif of the DREB/CBF family of transcription factors, in the promoters of selected genes. We found a single DRE/CRT motif in the promoters of MtCHS2, MtFLS, and MtGSTF7, while MtCHS, MtDFR1, MtANS, MtANR, and MtLAR genes lacked this motif in their promoters (Supplementary File S1). Additionally, single DRE/CRT motifs were identified in the promoters of MtKFB2 and MtKFB3, but not in the strongly regulated MtKFB1, suggesting an indirect regulation. Since most of the key late anthocyanin biosynthetic genes were markedly regulated by MtCBF4 despite lacking core CBF binding motifs in their promoters, we hypothesized that these genes might be indirectly regulated by MtCBF4. These genes are also known to be transcriptionally regulated by MtLAP1, MtTT8, MtMYB2, MtWD40, and others. Our analysis revealed significant changes in the expression of these regulatory genes, except MtWD40 (Fig. 1A, 1C, S2, S5). This prompted us to examine the presence of the core DRE motif in their promoters. A thorough analysis showed two DRE/CRT motifs in the promoters of only MtLAP1, located at -1803 bp and -2323 bp upstream of the transcription start site (TSS) (Fig. 2A, Supplementary File S1).

**Figure 2.**
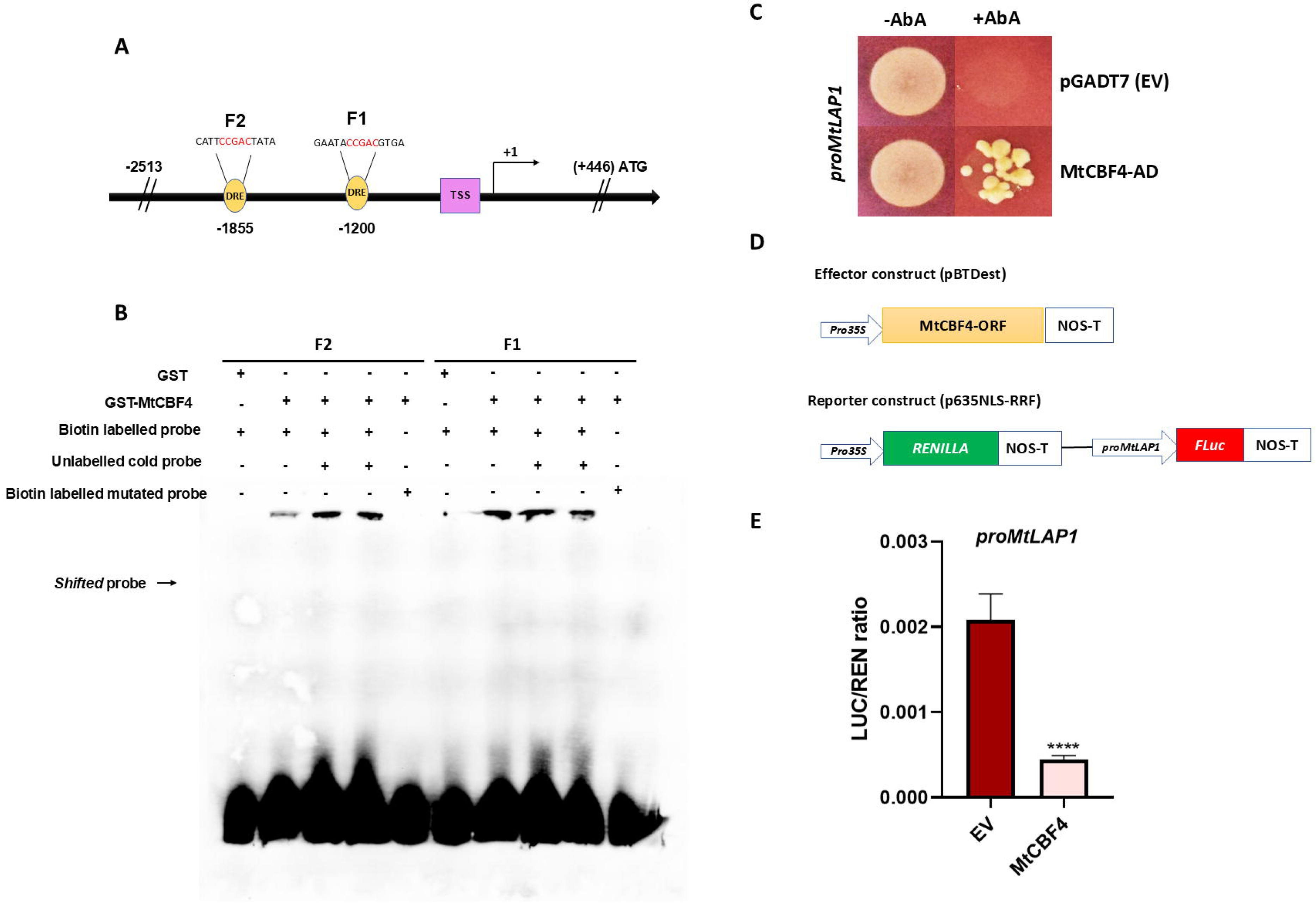
MtCBF4 binds to the *MtLAP1* promoter and represses its expression. **(A)** Illustration of the *cis*-motifs (DRE/CRT, RCCGAC) in the region located within 2513□bp upstream of the TSS in the promoter of *MtLAP1*. F1 and F2 fragments contain one DRE/CRT element each, having ACCGAC and CCCGAC motifs. **(B)** EMSA of MtCBF4 protein binding to promoter fragment F1 and F2 in vitro. A purified GST-MtCBF4 protein was incubated with biotin-labelled probes F1 and F2. The probes’ binding *cis*-motif and the mutated motif are highlighted in red in F1/F2 and F1m/F2m, respectively. + indicates presence and – indicates absence of probe/protein. **(C)** Yeast-one-hybrid (Y1H) assay showing the interaction of MtCBF4 and *MtLAP1* (614 bp) promoter fragment containing both the DRE/CRT elements (ACCGAC and CCCGAC). AD represents the empty pGADT7 vector. Growth was checked on synthetic-Leu dropout medium supplemented with AbA. **(D)** Schematic diagrams showing the reporter and effector constructs used for promoter-luciferase reporter assays. **(E)** The quantitative result of dual-luciferase reporter assays. The ratio of LUC and REN represents the repression efficiency of effectors (EV and MtCBF4) when co-transformed with the same *proMtLAP1-LUC* reporter. The data are mean values ±SD from three independent biological replicates.

To check whether MtCBF4 directly binds to the promoter of *MtLAP1*, we performed an Electrophoretic Mobility Shift Assay (EMSA) with two different probes, F1 and F2, each having two different DRE motifs of the *MtLAP1* promoter (Fig. 2A). A sharp shift was observed when purified GST-MtCBF4 was incubated with both the labelled probes, which substantially diminished when the mutated probe was added (Fig. 2B). This confirms the direct binding of MtCBF4 to the *MtLAP1* promoters *in vitro*. To further validate the direct binding, we isolated the 614 bp promoter fragment having both the DRE motifs and performed Yeast-One hybrid assay (Y1H). Y1H assay suggested transactivation of *MtLAP1* by binding of MtCBF4 in its promoter (Fig. 2C). As MtCBF4 down-regulated the expression of *MtLAP1* in *M. truncatula*, we sought to analyse the repression potential of the former over the promoter of the latter. To check whether MtCBF4 represses the promoter activity of *MtLAP1*, we isolated a 2513 bp promoter fragment upstream of TSS into the p635nRRF vector and performed dual-luciferase reporter assay with MtCBF4 as effector (Fig. 2D). MtCBF4 significantly repressed *MtLAP1* promoter activity in the dual luciferase reporter assay as indicated by reduced LUC/REN ratio (Fig. 2E). Together, these results suggest that MtCBF4 directly represses *MtLAP1* expression in *M. truncatula*.

### MtCBF4 directly activates MtJMJ13 expression to modulate histone H3K27 methylation

Transcription factors also influence chromatin state either *via* recruiting histone modifiers or regulating their expression (Candela-Ferre et al., 2024; Kim et al., 2024). Instances of such are the physical interaction of AtWRKY40-AtJMJ17, AtWRKY53-AtHDA9, AtBES1- AtREF6/ELF6, AtPIF7-AtREF6, and transcriptional up-regulation of AtREF6 by AtHSFA2 (Yu et al., 2008; Liu et al., 2019; Zheng et al., 2020; Wang et al., 2021; Cheng et al., 2024). In the transcriptome data of *MtCBF4-overexpressing* transgenic lines, we found two probable epigenetic regulators that were significantly up-regulated (Zhang et al., 2016). One of these up-regulated genes, *Medtr1g080340*, is a homolog of SUVR4 in Arabidopsis that encodes a Histone-lysine N-methyltransferase (H3K9me2 methyltransferase). Another gene, *Medtr5g029370*, a homolog of JMJ13 of Arabidopsis, encodes a lysine-specific demethylase (H3K27me3 demethylase). Among different epigenetic modifiers, Jumonji domain-containing demethylase has been widely implicated in abiotic stress response. Modifications like H3K27 and H3K4 trimethylations are the predominant methylation marks associated with cold stress response in plants (Liu et al., 2019; Zeng et al., 2019; Dasgupta et al., 2022; Faivre et al., 2024; Wang et al., 2024). Therefore, we hypothesized that Medtr5g029370 might be influencing chromatin state as a means of epigenetic regulation under cold stress downstream of MtCBF4 in *M. truncatula*. We named the gene *MtJMJ13* due to its highest sequence similarity (>70%) with AtJMJ13. Phylogenetic analysis showed clustering with AtJMJ13, but not with other characterized JMJs of Arabidopsis (Lopez et al., 2022). MtJMJ13 contained predicted Jumonji domain, a zinc finger domain, and N- and C-terminal flexible regions (Fig. S6). Sub-cellular localization in *Nicotiana benthamiana* leaf confirmed the predicted nuclear localization, which colocalized with a nuclear marker (Fig. 3A).

**Figure 3.**
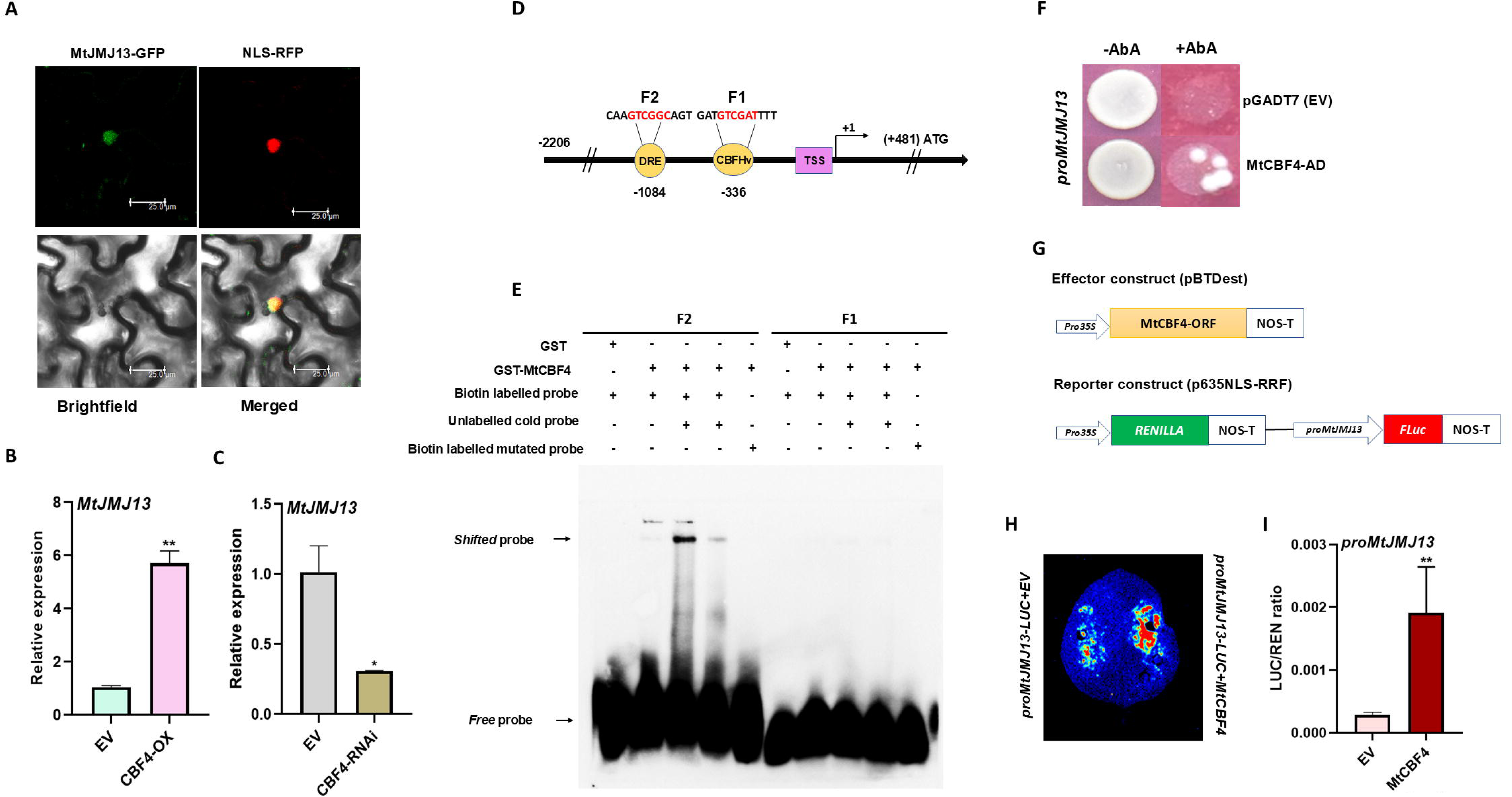
MtCBF4 promotes the expression of MtJMJ13 by directly binding to its promoter. **(A)** The subcellular localization of MtJMJ13 in *N. benthamiana* leaves epidermal cells. The subcellular location of free GFP protein (empty pUBICcGFP-DR) served as the negative control in confocal microscopy. NLS-RFP red fluorescence acted as a nuclear marker. Scale bar, 25□μm. **(B)** Relative expression level of *MtJMJ13* in *M. truncatula* leaves transiently expressing *MtCBF4* and the empty vector control, as determined by RT-qPCR. *MtACTIN* was used as the reference gene. **(C)** Relative expression level of *MtJMJ13* in *M. truncatula* hairy roots silencing *MtJMJ13* and the empty vector control, as determined by RT-qPCR. *MtACTIN* was used as the reference gene. **(D)** Illustration of the cis-motifs (DRE/CRT, RCCGAC; CBFHv, RYCGAC) in the region located within 2206□bp upstream of the TSS in the *MtJMJ13* promoter. F1 and F2 fragments each contain one DRE/CRT/CBFHv element *N. benthamiana* leaf image displays the original chemiluminescence picture after 150 s of exposure in EV and *MtCBF4* expressed in *N. benthamiana* leaves, along with the reporter. **(I)** The quantitative result of dual-luciferase reporter assays. The ratio of LUC and REN represents the promoter activation efficiency of effectors (EV and MtCBF4) when co-transformed with the same *proMtJMJ13-LUC* reporter. The data are mean values ±SD from three independent biological replicates.

To substantiate our hypothesis, we first checked the expression of *MtJMJ13* in *MtCBF4-*overexpressing leaves and *MtCBF4*-silenced hairy roots. In *MtCBF4- overexpressing* leaves, *MtJMJ13* transcript level increased by more than 5-fold, whereas *MtCBF4*-silenced hairy root tissues accumulated significantly decreased transcript level (Fig. 3B,3C). Further, promoter analysis of *MtJMJ13* detected the presence of a single DRE/CRT core motif at 1084 bp upstream of TSS, and a CBFHv motif (RYCGAC) at 336 bp upstream of TSS (Figure 3D). These findings prompted us to investigate the direct binding of the *MtJMJ13* promoter by MtCBF4. We designed probes containing both the DRE/CRT (F2) and CBFHv (F1) motifs and performed EMSA with purified GST-MtCBF4 (Fig. 3D). A sharp shift was observed when the DRE/CRT motif-containing probe was used, but not in the case of CBFHv probe (Fig. 3E). This confirms the direct binding of MtCBF4 to the promoter fragment F2 of *MtJMJ13 in vitro*. To further validate this interaction, we performed a Y1H assay with a 111 bp promoter fragment containing the DRE/CRT motif. Y1H suggested transactivation of 111 bp promoters of *MtJMJ13* by the binding of MtCBF4 *(Fig. 3F)*. Finally, we performed a dual luciferase assay to examine the transactivation potential of MtCBF4 *in planta* over the 2206 bp promoter fragment of *MtJMJ13 (Fig. 3G)*. Luciferase activity increased significantly when the *proMtJMJ13*-LUC reporter was co-infiltrated with MtCBF4 effector compared to EV (Fig. 3H). LUC/REN ratio increased significantly in the case of MtCBF4 co-expression compared to EV, suggesting its transactivation activity over *MtJMJ13* promoters (Fig. 3I). Taken together, these results conclude that MtCBF4 positively regulates the expression of *MtJMJ13* directly, and the latter might be playing roles downstream to the former in the cold stress signalling pathway in *M. truncatula*.

### MtJMJ13 represses anthocyanin biosynthesis

Previous reports have demonstrated histone methylation-mediated regulation of anthocyanin synthesis. For instance, JMJ25 and BONSAI Methylation 1 (IBM1), a JmjC domain-containing H3K9me2 demethylase, negatively regulate anthocyanin biosynthesis in poplar and Arabidopsis, respectively (Fan et al., 2018; Fan et al., 2024). We advantageously found a comparative transcriptome data set of the *jmj13* mutant and wildtype, and noted that many flavonoid biosynthesis genes, along with regulatory genes, were highly up-regulated in the *jmj13* mutant compared to wildtype in Arabidopsis (Yan et al., 2018). In contrast to this, *KFB^CHS^*, and *MYBL2* encoding genes were highly down-regulated. Interestingly, DREB1B/C/E/F genes were up-regulated and DREB2B/C/H genes were down-regulated (Yan et al., 2018; Zheng et al., 2019). We speculated that MtJMJ13 may also regulate the expression of anthocyanin biosynthetic genes in *M. truncatula*.

To investigate the role of MtJMJ13 in the regulation of anthocyanin biosynthesis, we transiently overexpressed *MtJMJ13* in the mature leaves of *M. truncatula* A17 ecotype. The expression of the *GFP* in the transformed leaves, as detected through semi-quantitative PCR, confirms successful transformation (Fig. S7). Subsequent expression analysis by RT-qPCR revealed more than 7-fold increase in the expression level of *MtJMJ13* in *MtJMJ13-* overexpressing leaves as compared to empty vector control (Fig. 4A). Further, expression analysis demonstrated decreased transcript levels of key biosynthetic genes like *MtCHS*, *MtDFR1*, along with the regulatory genes *MtLAP1, MtTT8,* and *MtWD40-1* (Fig. 4A, Fig. S8). A significant decrease in the expression of *MtGSTF7* was also observed (Fig. S8). We also analysed the expression of *MtCBF4* and found a ∼5-fold increase in its transcript level in *MtJMJ13* overexpressing leaves (Fig. 4A). Intriguingly, we observed up-regulation of *MtMYB2*, *MtKFB1*, and *MtKFB2* genes upon *MtJMJ13* overexpression in leaves (Fig. 4A, Fig. S8). Finally, we performed metabolite analysis through LC-MS. As expected, the content of major anthocyanidins like cyanidin, malvidin, peonidin, and petunidin decreased in the *MtJMJ13*-overexpressing leaves (Fig. 4B). This suggests that MtJMJ13 can repress anthocyanin biosynthesis in *M. truncatula*.

**Figure 4.**
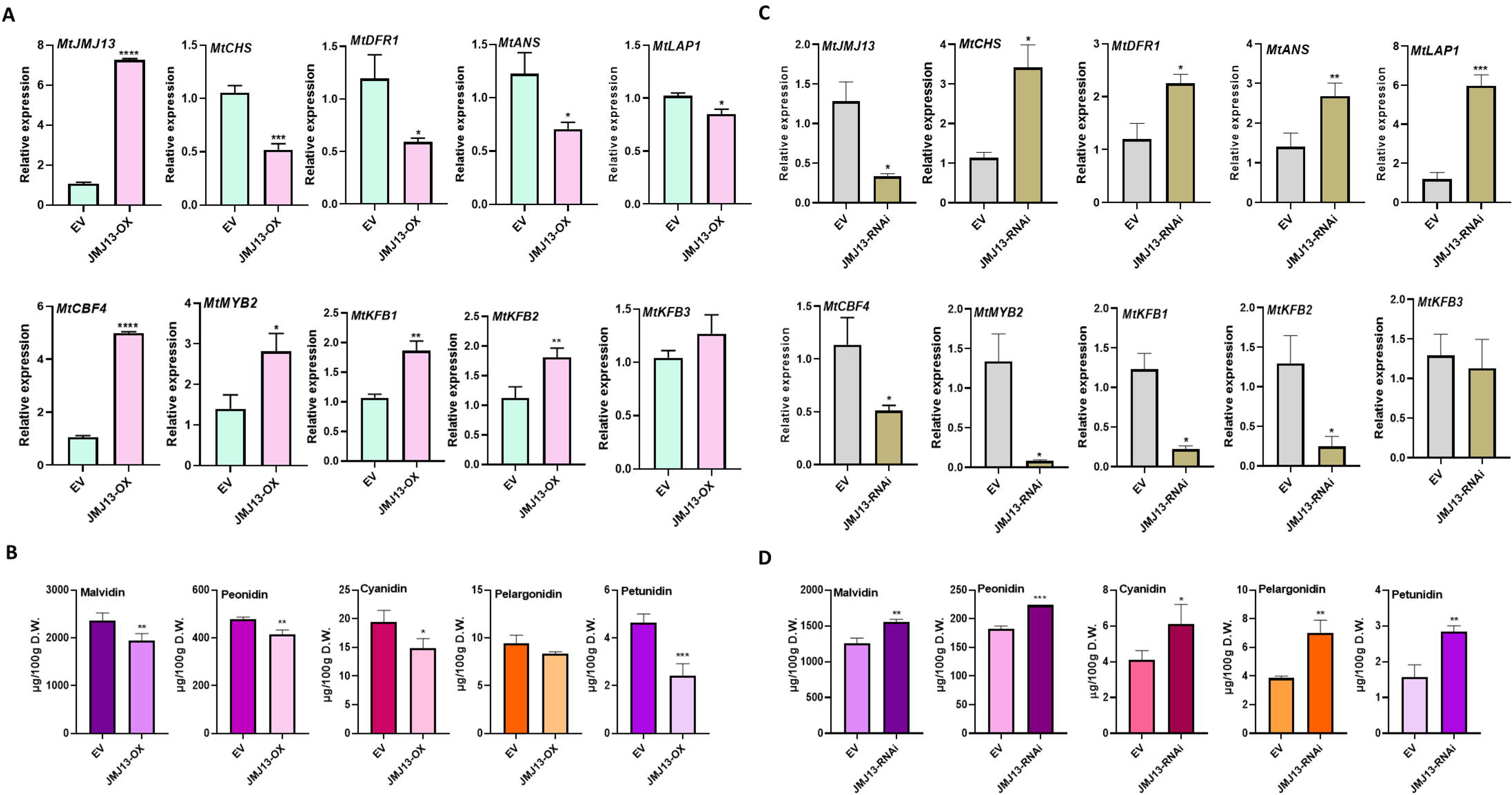
MtJMJ13 is a negative regulator of anthocyanin biosynthesis in *M. truncatula*. **(A)** Relative expression levels of *MtJMJ13*, biosynthesis genes (*MtCHS*, *MtDFR1*, *MtANS*), regulatory genes (*MtLAP1*, *MtCBF4*, *MtMYB2*, *MtKFB1/2/3*) in leaves that transiently overexpress *MtJMJ13*, as determined by RT–qPCR. *MtACTIN* was used as the internal control. **(B)** Quantitative targeted LC- MS-based quantification of major anthocyanidin (cyanidin, malvidin, peonidin, pelargonidin and petunidin) content in leaves transiently overexpressing *MtJMJ13*. The data are mean values ±SD (n=3 biological), each derived from several hairy roots. **(C)** Expression level of *MtJMJ13*, biosynthesis genes, and regulatory genes in transformed hairy roots silencing *MtJMJ13*. **(D)** Targeted LC–MS analysis of major anthocyanidins content in transformed hairy roots silencing *MtJMJ13*. The data are mean values ±SD (n=3 biological), each derived from several hairy roots.

For further validation, we performed gene silencing via RNAi and generated *MtJMJ13*-silenced hairy roots of *M. truncatula*. Bright red fluorescence of transformed hairy roots under a fluorescence microscope due to the expression of the marker gene DsRed confirmed successful transformation (Fig. S9). Further expression analysis *via* RT-qPCR revealed significantly decreased expression of *MtJMJ13* in the transgenic hairy roots (Fig. 4C). As anticipated, many anthocyanin biosynthetic genes like *MtCHS*, *MtCHI*, *MtDFR1*, *MtANS*, *MtLAR*, as well as the regulatory genes *MtLAP1*, *MtTT8*, displayed increased transcript levels in the *MtJMJ13*-silenced hairy roots (Fig. 4C, Fig. S10). Concomitantly, the expression level of *MtCBF4*, *MtMYB2*, *MtKFB1*, and *MtKFB2* decreased manifold, while no alteration in the transcript level of *MtWD40-1* and *MtKFB3* was found (Fig. 4C, Fig. S10). Lastly, metabolite analysis showed substantially increased accumulation of all the major forms of anthocyanidins, (Fig. 4D). Altogether, we concluded that MtJMJ13 is a negative regulator of anthocyanin synthesis in *M. truncatula*.

### MtJMJ13 directly demethylates the gene body of *MtCBF4* to activate its expression

AtJMJ13 is known to recognise H3K27me3 peptide, and specifically demethylate H3K27me3, but not the other histone methyl marks in Arabidopsis (Zheng et al., 2019). MtJMJ13 shares more than 70% protein sequence similarity with AtJMJ13, and the catalytic domain has high sequence conservation (Fig. S6). We checked the level of H3K27me3 in *MtJMJ13*-silenced hairy roots, which displayed increased H3K27me3 levels compared to the control. This suggests that MtJMJ13 is also a H3K27me3 demethylase (Fig. S11).

Intriguingly, in our study, only the negative regulators of anthocyanin biosynthesis, *viz. MtCBF4, MtMYB2,* and *MtKFB1-3* were found to be positively regulated by MtJMJ13. Further inspection into the comparative ChIP-seq data of wildtype and *ref6-elf6^C^ jmj13^G^* triple mutant in Arabidopsis disclosed increased H3K27 trimethylation in the entire gene body of *MYBL2*, *KFB^CHS^*, and *DREB2* genes (Yan et al., 2018). Therefore, we hypothesized that MtJMJ13 might directly associate with and demethylate the gene bodies of *MtCBF4*, *MtMYB2*, *MtKFB1*, and *MtKFB2*. To analyse the H3K27me3 methylation status of these genes in *MtJMJ13*-silenced hairy root tissues, we performed ChIP-qPCR. A series of primers was designed from the different regions of their gene body, and the antibody specific to H3K27me3 was used for immunoprecipitating the DNA-protein complex (Fig. 5A-D). Methylation level increased significantly in the F3 region of *MtCBF4*, whereas no significant changes were detected in other regions (Fig. 5A). On the other hand, the *MtKFB1* and *MtKFB2* gene bodies had higher methylation levels at the F1 and/or F2 regions in their gene body, closer to the TSS (Fig. 5B-C). In the case of *MtMYB2*, a significant increase in the methylation level was found at F2 and F4 regions (Fig. 5D). Hypermethylation of H3K27 in these regions demonstrated that MtJMJ13 can dynamically demethylate the gene body of these target genes, like *MtCBF4*, *MtMYB2*, and *MtKFB1/2*, leading to a transcriptionally active chromatin state and enhanced gene expression.

**Figure 5.**
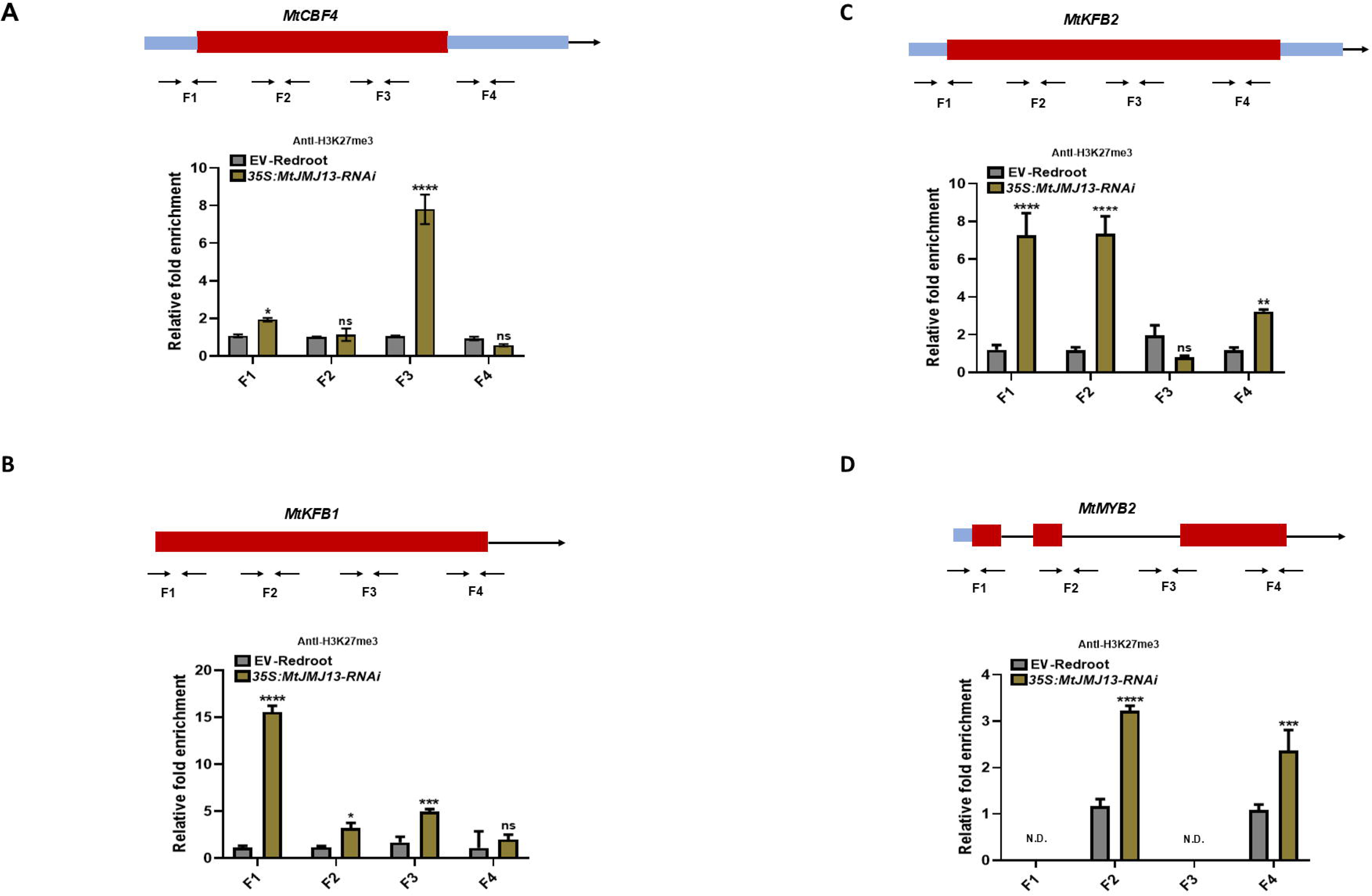
MtJMJ13 targets specific regions and demethylates *MtCBF4*, *MtKFB1*, *MtKFB2*, and *MtMYB2* chromatin in *M. truncatula*. Schematic genetic structures of various genes and the regions used for ChIP-qPCR. The burgundy red boxes represent exons; the black boxes represent 5′ and 3′ UTRs; the black lines represent introns or intergenic regions. Gene body structure and detection of the H3K27me3 level in **(A)** *MtCBF4*, **(B)** *MtKFB1*, **(C)** *MtKFB2*, and **(D)** *MtMYB2* chromatin by ChIP-qPCR. The results were calculated as relative fold enrichment as compared to the control, which was set to 1. Data is represented as mean ± SD of three technical replicates of one representative experiment; the experiment was repeated twice with similar results.

### Cold stress induces anthocyanin biosynthesis in *M. truncatula* seedlings

MtCBF4 is known to be involved in freezing tolerance in *M. truncatula* (Li et al., 2011; Zhang et al., 2016). Cold or freezing stress leads to both up-regulation and down-regulation of anthocyanin biosynthesis in different plant species (An et al., 2020; Mao et al., 2022; Jiang et al., 2022). To explore the regulation of anthocyanin biosynthesis by cold stress signalling pathway in *M. truncatula*, four-week-old plants were subjected to low temperature conditions and kept at 4L, while control plants were kept at 22L. Time course metabolite analysis of the extract was performed at different time points of cold treatment (0, 1, 3, 6, 12, 24, 48 hrs) (Fig. 6A). Total anthocyanin content analysis showed increased anthocyanin accumulation upon cold treatment, which started decreasing after 12 hrs (Fig. 6B). LC-MS analysis further confirmed increased accumulation of various anthocyanidins like cyanidin, delphinidin, malvidin, petunidin and peonidin (Fig. 6C). These results suggest that cold stress induces greater anthocyanidin accumulation in *M. truncatula*.

**Figure 6.**
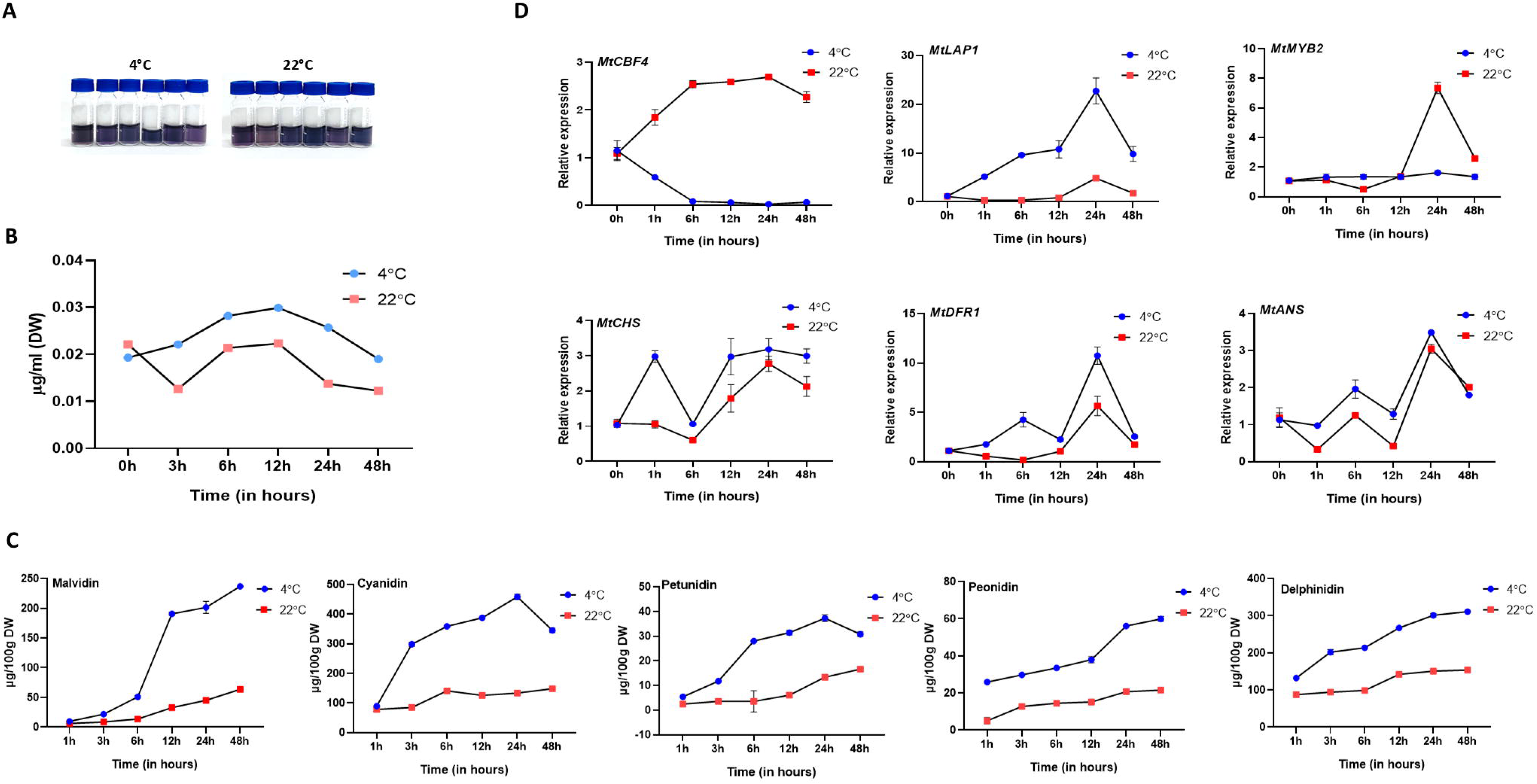
Cold stress induced anthocyanin synthesis in the leaves of *M. truncatula* A17 ecotype. **(A)** Methanolic extracts of 2-month-old *M. truncatula* plants exposed to either cold stress (4□) or optimal growth temperature (22□) as a control. **(B)** Total anthocyanin content in the extracts of cold and control-treated plants at the indicated time points (1 h, 3 h, 6h, 12h, 24h, 48h). **(C)** Targeted LC– MS analysis of major anthocyanidins (cyanidin, malvidin, peonidin, pelargonidin and petunidin) content in cold vs control plants. The data are mean values ±SD (n=3 biological). **(D)** Relative expression of *MtCBF4*, *MtLAP1, MtMYB2,* and their downstream biosynthetic genes (*MtCHS, MtDFR1,* and *MtANS*) in the leaves of cold vs control plants at different time points. The values were normalized to *MtACTIN* expression. The data represent the mean ± SD of 3 biological replicates.

Further, we carried out expression analysis via RT-qPCR, which suggested differential regulation of biosynthesis and regulatory genes. Many biosynthesis genes like *MtPAL*, *MtCHS*, *MtF3H*, *MtF3’H*, *MtDFR1*, *MtANS*, and *MtGSTF7* were analysed at different time points of cold treatment (Fig. 6D, Fig. S12). Interestingly, *MtCHS, MtDFR1*, and *MtANS*, the key genes for anthocyanin synthesis, exhibited increased transcript accumulation up to 24 hours of treatment (Fig. 6D). Similarly, the expression level of *MtLAP1*, *MtTT8*, and *MtWD40-1* also increased along with their target genes (Fig. 6D, Fig. S12). Surprisingly, we observed a substantial decrease in the transcript level of *MtCBF4* under cold stress (Fig. 6D), which corroborated with a previous study, where the transcript level increased at the initial time points, and started decreasing after 3.5 hrs of cold treatment and remained lower up to 24 hrs (Zhang et al., 2016). On the other hand, the expression level of *MtJMJ13*, *MtMYB2*, *MtKFB1*, and *MtKFB2* didn’t display significant up-regulation (Fig. S12). Taken together, we concluded that cold stress induces the accumulation of anthocyanin *via* up-regulation of anthocyanin biosynthesis-related genes while suppressing the expression of *MtCBF4*, the repressor of anthocyanin biosynthesis.

## Discussion

Molecular intricacies governing abiotic stress response in plants have been meticulously studied in recent years, which have greatly advanced our understanding of plant stress physiology. Low temperature has been previously reported to both promote and repress anthocyanin biosynthesis in plants *via* switching on the downstream signal transduction pathway (Ahmed et al., 2015; Schulz et al., 2025; Wang et al., 2024; Lin et al., 2025). Anthocyanins, in turn, protect the plants from oxidative damage and membrane lipids from peroxidation during stress conditions (Mahmoud et al., 2024; Liu et al., 2025). In this study, we explored a novel regulatory loop that functions to inhibit anthocyanin biosynthesis in *M. truncatula* (Fig. 7). We report that MtCBF4 represses anthocyanin accumulation *via* both down-regulation of anthocyanin biosynthetic genes and up-regulation of negative regulator encoding genes. MtCBF4 directly binds and represses the expression of *MtLAP1,* while directly activating *MtJMJ13*. MtJMJ13 in turn activates the expression of *MtCBF4 via* chromatin histone H3K27me3 demethylation, thus forming a positive feedback loop (Fig. 7). This positive feedback loop seems to be partly suppressed under cold stress, as evident from the down-regulation of *MtCBF4* and anthocyanin biosynthesis genes under prolonged cold treatment. Thus, cold promotes anthocyanin accumulation in *M. truncatula* (Fig. 7). Our results show that low temperature treatment downregulates the expression of *MtCBF4*, which acts as a suppressor of *MtLAP1* expression and anthocyanin biosynthesis. On the other hand, MtCBF4 also acts as a positive regulator of a chromatin demethylating enzyme MtJMJ13. Under low temperature, downregulation of MtCBF4 would result in decreased expression of MtJMJ13, leading to increased H3K27 methylation of the gene body of *MKFB1*, *MtKFB2*, *MtMYB2*, and *MtCBF4*, and subsequently activation of anthocyanin biosynthetic genes.

**Figure 7.**
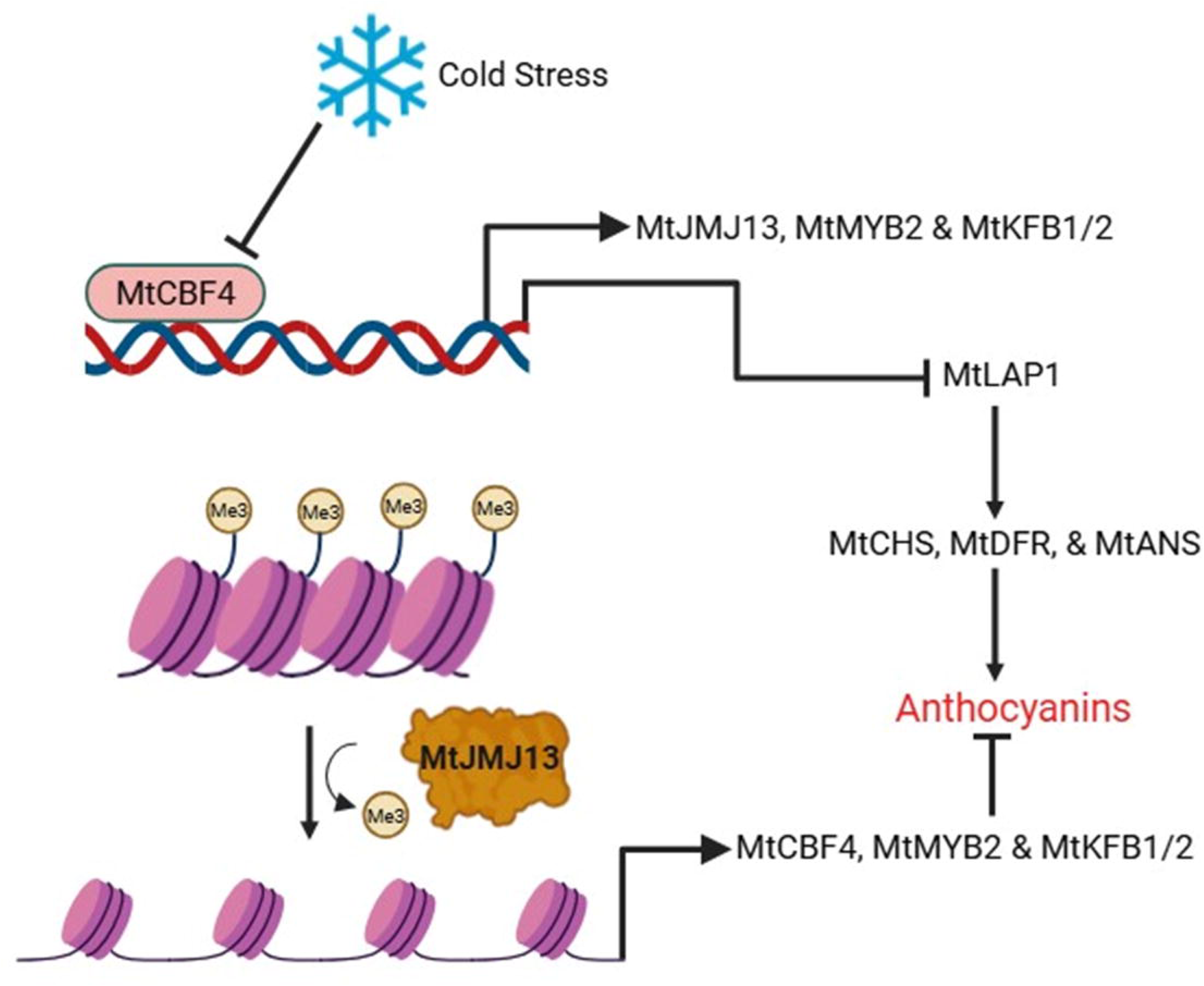
The working model of MtCBF4-MtJMJ13 positive feedback loop controlling anthocyanin biosynthesis in *M. truncatula*. MtCBF4 directly binds to the promoter and represses the expression of *MtLAP1*, leading to the down-regulation of downstream biosynthesis genes and ultimately, decreased anthocyanin accumulation. Cold stress represses the expression of *MtCBF4* to promote anthocyanin accumulation. *MtCBF4* also regulates the expression of *MtJMJ13* by binding to its promoter and enhancing its expression. *MtJMJ13*, in turn, precisely targets specific regions of the *MtCBF4* gene body and demethylates H3K27me3, leading to the transcriptional activation of the latter. It also demethylates *MtKFB1*, *MtKFB2*, and *MtMYB2*. All these events lead to increased expression of *MtJMJ13* target genes and decreased expression of *MtLAP1* and other biosynthesis genes. Thus, in this way, MtJMJ13 acts as a negative regulator of anthocyanin biosynthesis. The MtCBF4-MtJMJ13 positive feedback regulatory loop thus operates to control anthocyanin biosynthesis, and cold partly attenuates the loop via down-regulation of *MtLAP1*, which might lead to increased anthocyanin accumulation under cold stress in *M. truncatula*.

### MtCBF4 as a key suppressor of anthocyanin accumulation in *M. truncatula*

As previously mentioned, MtCBF4 is a known regulator of salt and freezing stress tolerance in *M. truncatula* (Li et al., 2011; Zhang et al., 2016). In Arabidopsis, it regulates COR15A, COR15B, KIN1, RD17, RD29A, and RD29B, the downstream targets of AtDREBs (Li et al., 2011). In our study, we found that cold stress reduced the transcript level of *MtCBF4* (Fig. 6D), which is quite consistent with a previous study, where the *MtCBF4* transcript level decreased after a few hours of cold treatment (Zhang et al., 2016). In our study, we found that anthocyanin content increased and was negatively correlated with the expression of *MtCBF4* upon cold treatment (Fig. 6C-D). Interestingly, *MtCBF2* and *MtCBF3* were found to be located in a major freezing tolerance QTL region of *M. truncatula* chromosome 6 (Tayeh et al., 2013), but not *MtCBF4*, which is present on chromosome 1 (Zhao et al., 2023). MtCBF4, however, regulates many downstream COR genes, like *MtCAS15*, *MtGolS1*, and *MtGolS2*, leading to raffinose accumulation and cold acclimation (Sun et al., 2021). MtCBF4 strongly down-regulates many genes involved in the biosynthesis and regulation of anthocyanin biosynthesis in *M. truncatula*. MtLAP1, an Arabidopsis PAP1 homolog, has been identified earlier to be a strong regulator of anthocyanin accumulation in *M. truncatula* and *M. sativa* (Peel et al., 2009). We found that MtCBF4 directly binds and represses *MtLAP1*. Many other biosynthetic genes were thus down-regulated, while negative regulators of flavonoid synthesis like *MtMYB2*, and the putative *MtKFB^CHS^* (*MtKFB1/2*) genes displayed up- regulation. These observations suggest a key role of MtCBF4 as the suppressor of anthocyanin accumulation in *M. truncatula*.

*MtLAP1* overexpression in *M. truncatula* and *M. sativa* not only increased anthocyanin biosynthetic genes like *MtPAL*, *MtCHS*, *MtCHI*, *MtF3’H*, *MtDFR1*, *MtANS*, but also genes that are involved in B-ring modification (*UGT78G1*, malonyltransferase), pathway regulation (*TTG2*, *AN1*), and transport (LeOPT1, MATE, GST) (Peel et al., 2009). It also suggests that the down-regulation of the expression of the *MtGSTF7* gene is MtLAP1- dependent, and MtCBF4 can also regulate other modifying and transport-related genes. Many such genes are also regulated by MtTT8, MtWD40, and MtMYB2, and up- or down- regulation of these genes by MtCBF4 might have resulted in the decreased transcript accumulation of biosynthesis genes (Pang et al., 2009; Jun et al., 2015; Li et al., 2016). *MtTT8* expression remarkably decreased in *MtCBF4* overexpressing tissues and increased in the silenced hairy roots. The absence of the DRE/CRT promoter hints at indirect regulation *via* some other regulators. Prior to our study, a few studies have demonstrated the regulatory role of DREB/CBF transcription factors in flavonoid biosynthesis. The UDP-glycosyltransferases, UGT79B2 and UGT79B3, catalyze anthocyanin rhamnosylation and are directly activated by AtCBF1 in response to low temperature in Arabidopsis (Li et al., 2017). CBFs also act synergistically with the MYB-bHLH-WD40 complex *via* direct interactions. For instance, SmCBF1/2/3 physically interacts with SmCBF113 to promote anthocyanin synthesis in eggplant (Zhou et al., 2020). Cold temperature also activates LcDREB2C, which in turn, directly targets many genes like *LcMYB1*, *LcCHI*, and *LcF3H* to enhance anthocyanin accumulation during fruit ripening in litchi (Zou et al., 2024). In our present study, we showed that MtCBF4 acts as a negative regulator of anthocyanin accumulation. Higher expression of *MtCBF4* in the root, particularly the tip region, than in other organs suggests a critical role of MtCBF4 in salt stress response However, leaves and stems accumulate higher transcript levels of *MtCBF1* and *MtCBF2* (Li et al., 2011). Anthocyanin accumulates mostly in shoots, and MtCBF4-mediated repression might be a functional module to suppress anthocyanin biosynthesis and divert the flux towards other metabolic pathways in *M. truncatula* under normal growth conditions. Interestingly, MtCBF4 forms a clade with DREB1-type sub-class, but shares the highest sequence similarity (57 %) to AtCBF4/AtDREB1D, which is known to be induced by salt, drought, and ABA (Li et al., 2011). As expected, it possesses a conserved AP2 DNA binding domain and a C-terminal LWSY motif, along with CBF characteristic sequence features (PKK/RPAGRxKFxETRHP, DSAWR, and A(A/V)xxA(A/V)xxF). However, it has a DSAWK motif instead of DSAWR (Li et al., 2011). Thus, it further confirms the conservation of structural and regulatory aspects of CBF/DREBs under abiotic stress in plants.

### MtJMJ13 as the downstream node in the MtCBF4-mediated signalling pathway

*M. truncatula* genome encodes 34 putative Jumonji domain-containing genes as compared to 20 that are present in *Arabidopsis thaliana* (Lopez et al., 2022). MtJMJ13 has been previously identified and named as MtJMJ25, which forms a cluster together with AtJMJ13 and thus belongs to the KMD4 clade (Lopez et al., 2022). It has conserved JmjN, JmjC, and C5HC2 motifs, and shares high sequence similarity to AtJMJ13, suggesting conservation of catalytic functions (Fig. S4). This gene family has been studied extensively in many plant species to regulate diverse aspects of plant development and stress response, as mentioned previously. However, the functional data of the gene family members in *Medicago sp.* remains largely unexplored. In our study, we identified a novel histone demethylase, which up-regulates diverse classes of regulatory genes known to suppress or might suppress anthocyanin accumulation, such as transcription factors (MtCBF4, MtMYB2) and F-box proteins (MtKFB1/2/3). It leads to MtCBF4-mediated repression of *MtLAP1*, and subsequently downregulation of anthocyanin biosynthesis genes. Apart from *MtLAP1, MtTT8* expression also decreased dramatically upon *MtJMJ13* overexpression. The expression of *AtTT8* is directly regulated by AtMYBL2 as well as MYB-bHLH-WD40 complex in a feedback loop (Baudry et al., 2006; Matsui et al., 2008). In this way, the decreased expression of *MtTT8* upon *MtJMJ13* overexpression might have resulted from the down-regulation of *MtLAP1* and up-regulation of *MtMYB2*. Thus, down-regulation of anthocyanin accumulation by MtJMJ13 is coordinated by the interplay of various regulatory factors acting downstream of MtJMJ13.

Prior to our study, a limited number of studies have shed light on the histone demethylase-dependent epigenetic regulation of anthocyanin biosynthesis. As an example, JMJ25 enhanced the expression of *MYB182*, but not *MYB57*, the two repressors of anthocyanin biosynthesis. Neither, it regulated the expression of *MYB116*, *MYB117*, *MYB118*, *MYB119*, and *MYB120*, known to promote anthocyanin synthesis, nor *MYB57* (Fan et al., 2018). JMJ25 directly associates with the gene body of *MYB182* and demethylates the chromatin, resulting in the higher expression of *MYB182*, and subsequently decreased expression of *DFRs*, *ANSs*, and *UFGTs* (Fan et al., 2018). Likewise, in our study as well, we discovered that MtJMJ13 demethylates the trimethylated H3K27 on the gene body of *MtCBF4* and *MtMYB2*, leading to their up-regulation (Fig. 5A, D). Eventually, *MtLAP1* and other biosynthetic genes were down-regulated. Thus, it can be stated that MtJMJ13 acts downstream of MtCBF4 to modulate anthocyanin biosynthesis *via* epigenetic regulation. In another study, high light induced BONSAI Methylation 1 (IBM1) strongly promotes H3K9me2 demethylation and non-CG DNA demethylation at the gene body of SPA1/3/4 (SUPPRESSOR OF PHYTOCHROME A), which further leads to higher expression of SPA genes and decreased anthocyanin accumulation of anthocyanin (Fan et al., 2024).

In an earlier study on tomato, it was reported that SlJMJ7, an H3K4 demethylase, represses the expression of DNA demethylase (*DML2*/*DEMETER*-like), therefore affecting global DNA methylation (Ding et al., 2022). JMJs are also known to interact with and recruit other members of the JMJ family (Chai et al., 2024). So, it is also possible that MtJMJ13 recruits other JMJs or chromatin modellers to modulate various histone methylations and also DNA demethylases at the target gene loci for DNA hypomethylation in *M. truncatula*. A thorough investigation is needed to unravel how MtJMJ13 may influence DNA methylation in *M. truncatula*.

### The MtCBF4-MtJMJ13 positive feedback loop might be conserved in plants

Surprisingly, we found that MtJMJ13 can demethylate the H3K27 trimethylated marks on the gene body of *MtCBF4*, thus forming a novel positive feedback loop strongly controlling anthocyanin accumulation in *M. truncatula*. In the comparative RNA-seq data of Arabidopsis *jmj13* mutant vs WT, we observed down-regulation of DREB2 genes, which could be successfully extrapolated to our study, although MtCBF4 doesn’t belong to the DREB2-type subfamily (Li et al., 2011; Zheng et al., 2019). This kind of feedback loop, consisting of a transcription regulator and a histone demethylase, has also been discovered earlier in Arabidopsis (Liu et al., 2019). Under heat stress in Arabidopsis, HSFA2 directly binds to the promoter and activates the expression of *REF6*, an H3K27me3 demethylase. REF6, in turn, demethylates HSFA2 and promotes its expression, thus forming a positive feedback loop (Liu et al., 2019). In our study, we discovered a MtCBF4-MtJMJ13 module as a novel positive feedback loop that might be conserved in land plants (Fig. 7).

Plant-specific KMD4 subfamily was first found in land plants but not in green algae (Lu et al., 2008; Qian et al., 2015). MtJMJ13, being a member of this subfamily, might have a conserved regulatory function like its orthologs in Arabidopsis. JMJ13 has been identified as the photoperiod and temperature-dependent regulator of flowering in Arabidopsis. The protein level of JMJ13 increases under heat, which may be lower under normal or low temperature conditions (Zheng et al., 2019). JMJ13 probably acts as the positive regulator of the floral repressor gene SHORT VEGETATIVE PHASE (*SVP*), and the early flowering phenotype of the *jmj13* mutant is due to lower expression of *SVP* (Yamaguchi., 2021). Thus, MtJMJ13 may also function as the repressor of flowering in *M. truncatula* in a temperature-dependent manner. JMJ13 is regulated at the protein level under high temperature in Arabidopsis; likewise, MtJMJ13 might also be regulated at the protein level apart from transcriptional regulation of MtCBF4. We speculate that MtCBF4 may also affect flowering at the upstream of *MtJMJ13*. DREB1A, DREB2C have been reported earlier to delay flowering in Arabidopsis (Suo et al., 2016; Song et al., 2020). MsCBF9 from *M. sativa* delayed flowering when overexpressed in Arabidopsis (Cui et al., 2025). Thus, it raises a greater possibility that MtCBF4 might also negatively influence flowering by affecting the expression of *MtJMJ13*. *MtCBF4* expression decreases under cold stress, further strengthening our hypothesis that cold-repressed MtCBF4 might have positive effects on cold-induced flowering in *M. truncatula*. Similar phenomena have also been reported earlier in litchi, where the expression of *LcCBF1/2/3* decreases under cold conditions, and it delays flowering in Arabidopsis (Shan et al., 2023).

Secondly, many flavonoid biosynthesis genes were up-regulated in the *jmj13* mutant of Arabidopsis, with decreased expression of *DREB2*, *MYB2*, and *KFB^CHS^* (Yan et al., 2018; Zheng et al., 2019). In our study as well, we noted that MtJMJ13 can directly and positively regulate the latter set of genes and indirectly inhibit the expression of anthocyanin biosynthesis genes. This suggests that the JMJ13 function might be conserved in land plants, or at least in dicots. *JMJ13* expression and protein level increase under heat conditions is known to suppress anthocyanin accumulation, again confirming the key role of JMJ13 under high temperature conditions (Zheng et al., 2019; Zhao et al., 2023).

### Cold stress attenuates the regulatory loop

In our study, cold temperature could curb the regulatory feedback loop, at least partly by decreasing the expression of *MtCBF4 (Fig. 6D)*. Thus, many anthocyanin biosynthetic genes are freed from suppression, leading to increased anthocyanin accumulation and greater cold/freezing tolerance. The positive role of cold stress in anthocyanin induction has been extensively studied in many crop plants (Mao et al., 2022; Jiang et al., 2022; Schulz et al., 2025; Wang et al., 2024; Lin et al., 2025). MYB30-INTERACTING E3 LIGASE 1 (MdMIEL1), an E3 ubiquitin ligase, negatively regulates anthocyanin biosynthesis in apple (An et al., 2017; An et al., 2020; An et al., 2021). The promoter activity, transcript level, and protein level of MdMIEL1 are reduced under cold stress in apple, which leads to increased anthocyanin accumulation (An et al., 2019). Likewise, the reduced transcript level of *MtCBF4* under cold stress might lead to increased anthocyanin accumulation under cold in *M. truncatula*.

In our study, we identified a new regulatory loop, which acts as a strong negative regulator of anthocyanin biosynthesis (Fig. 7). The possibility that it can also affect other traits like flowering, nodulation, and other pathways cannot be ruled out. Although many studies have shed light on the regulatory mechanisms of stress tolerance in *M. truncatula* and *M. sativa*. Here, we proposed a working model that can be effectively used for genetic engineering to improve the mentioned traits in an important forage crop like Medicago.

## Materials and methods

### Plant materials and growth conditions

*Medicago truncatula* ecotype A17 was used for various experiments in this study. Seeds were treated with concentrated sulfuric acid for 5 min, and then washed five times with cold sterile water. Acid-treated seeds were sterilized with 30% bleach (sodium hypochlorite) for 5 min, followed by washing three times with cold sterile water. Sterilized seeds were stratified at 4°C overnight and germinated on filter paper in petri plates at 22°C in the dark before transferring to soil. The plants were grown in a growth chamber (Percival AR-41 L3, Perry, IA, USA) at 22°C under 45 % RH and a 16/8 hours day/night cycle at 100 mmol m^−2^ s^−1^.

*Nicotiana benthamiana* was germinated and grown on plates containing Murashige and Skoog medium at 24°C under a 16/8 hours photoperiod at 100 mmol m^−2^ s^−1^ for 1 week, then transplanted to a soil: vermiculite (1:1, v/v) mixture and grown in a greenhouse at 22°C under a 16-h photoperiod for 5 weeks.

The cold treatment assay for *M. truncatula* was conducted as described in previous studies (Zhao et al., 2020; Ye et al, 2024). Briefly, 2-month-old A17 ecotype plants grown in a soil: vermiculite (1:2, v/v) were used for the cold treatment assay. For treatment, some plants were kept in a growth chamber under optimal growth conditions, except the temperature, which was set to 4°C. Control plants were kept at 22°C. Leaf samples were harvested at different time points and frozen in liquid N_2_ for further use.

### Phylogenetic analysis and sequence alignment

To identify the KFB^CHS^ gene family, the protein sequence of AtKFB^CHS^ and Medtr5g033880 were used for pBLAST against the *M. truncatula* A17 reference genome (JCVI-Mt. 4.0) (https://medicago.toulouse.inra.fr/MtrunA17r5.0-ANR/). Protein sequences of MtKFB1-3, AtKFB^CHS^, and other characterized KFB genes were used for constructing a phylogenetic tree using MEGA-X software via the maximum-likelihood method with 1000 bootstrap values (Kumar et al., 2018). Multiple sequence alignment was performed on MAFFT v.7.299b (Katoh & Standley, 2013).

The protein sequences of JMJs characterized in Arabidopsis and other plants were retrieved from NCBI, and together with MtJMJ13, were used in phylogenetic analysis using MEGA X via the maximum-likelihood method with 1000 bootstrapping rounds. For multiple sequence alignment, the MtJMJ13 peptide sequence was aligned with other JMJs characterized in Arabidopsis and other plants using MAFFT v.7.299b (Katoh & Standley, 2013). Different domains and motifs were identified manually after comparison with their functional orthologs.

### RNA isolation, first-strand cDNA synthesis, and RT-qPCR analysis

Total RNA was isolated from different *M. truncatula* samples using the Spectrum™ Plant Total RNA Kit (Sigma-Aldrich) and treated with RNase-free DNase (TURBO DNA-free™ Kit, Invitrogen). First-strand cDNA was generated from total RNA using a RevertAid H Minus First Strand cDNA Synthesis Kit (Thermo Fisher) and oligo(dT) primers. RT-qPCR-based gene expression analyses were carried out on a 7500 Fast Real-time PCR System (Applied Biosystems) using the 2x SYBR Green PCR Master mix (Applied Biosystems) and diluted cDNA in a final volume of 10 µL. *MtACTIN* (Naik et al., 2021) was used to normalize transcript abundance. Relative expression levels were calculated via the cycle threshold (C_t_) 2^−ΔΔCT^ method (Livak & Schmittgen,L2001), and are presented as fold changes relative to the control condition, depending on the conditions. Three biological replicates were analysed in each RT-qPCR analysis as specified.

### Cloning of full-length CDS from cDNA of *M. truncatula*

The full-length CDSs (without the stop codons) of *MtCBF4* and *MtJMJ13* were amplified using first-strand cDNA of *M. truncatula* A17 ecotype leaves and integrated into pENTR™/D-TOPO™ (Invitrogen). Further, Sanger sequencing of the resulting entry plasmids was performed by the NIPGR sequencing core facility (New Delhi, India). The set of primers used for the amplification of CDS is provided in Supplementary File S2.

### Subcellular localization and co-localization assays

CDS of *MtJMJ13* was cloned into the pUBIcGFP-DR binary vector (Chakrabarty *et al*., 2007), generating a C-terminal green fluorescent protein (GFP) fusion protein construct. An empty vector was used as a negative control. The plasmid was transformed into *Agrobacterium tumefaciens* strain GV3101-pMP90 (Koncz & Schell,L1986). Equal volumes of *A. tumefaciens* cultures having the desired gene and the NLS-RFP marker were mixed for agro-infiltration into the abaxial side of *N. benthamiana* leaves, according to the protocol described by Walter *et al*. (2004). Infiltrated plants were grown for 2 days in the dark at 25°C. Leaf discs were cut from the infiltrated leaves and observed under an argon laser confocal laser-scanning microscope (TCS SP5; Leica Microsystems, Wetzlar, Germany) with YFP and RFP filters. YFP and RFP fluorescence was observed at 514-nm excitation/527-nm emission and 558-nm excitation/583-nm emission wavelengths, respectively.

### Transient expression in adult leaves

Transient expression in leaves was performed as described previously (Picard et al., 2013). The coding sequence of *MtCBF4* and *MtJMJ13* was subcloned into *pUBIcGFP-DR* (Kryvoruchko et al., 2016) vector to create the *pZmUBI:MtCBF4-GFP* and *pZmUBI:MtJMJ13-GFP* constructs, and subsequently integrated into *A. tumefaciens* strain EHA105. The transformed *A. tumefaciens* were grown at 28°C on a shaker overnight. The cultures were centrifuged at 4500 ×*g* for 10 min, followed by resuspension of the pellet in infiltration buffer (10 mM NaCl, 1.75 mM CaCl_2_, 2 µl Tween-20, 100 mM acetosyringone) to an OD_600_ of 0.5. The fully unfolded trifoliate healthy leaves of 3–4-week-old *M. truncatula* A17 plants were selected for infiltration using a syringe. Plants were kept in the dark at 22 °C overnight and transferred to the optimum growth conditions at 22°C. Infiltrated leaf samples were harvested after 2-3 days and frozen in liquid N_2_ for further use.

### Hairy root transformation of *M. truncatula*

For gene silencing via hairy root transformation, we utilized the 250-350 bp CDS-3’ UTR region of *MtCBF4* and *MtJMJ13* in *pK7GWIWG2(II)-RedRoot* and empty vector control (Sinharoy et al., 2015). The *M. truncatula* accession A17 was used for hairy root transformation with the *Agrobacterium rhizogenes* ARqua1 strain as described previously (Boisson-Dernier et al. 2001). Sterilized seeds were kept in the dark till germination. Seedlings with approx. 1 cm long roots were cut at the middle of the root, and the cut end was coated with ARqua1 culture by gently scraping. Finally, the seedlings were kept and placed on Fahraeus Media and incubated at 22°C with 14–10 h light–dark photoperiods for 12–14 days. The transformed hairy roots were identified under a stereomicroscope, showing RFP fluorescence using a red filter/RFP filter.

### Anthocyanin content analysis

Total anthocyanin content (TAC) was measured in the sample as described previously (Pal et al., 2022) with minor modifications. Briefly, 2 ml of acidified methanol (1% v/v) was mixed with 0.25 g of homogenized tissue, followed by incubation at 28°C overnight in the dark. The extract was centrifuged at 10000g, absorbance of the supernatant was measured at 530 and 657 nm. The formula for calculating TAC is (A530–0.25*A657)/weight of tissue (g) (Rajput et al., 2025).

### LC-MS analysis of anthocyanin quantification

Anthocyanidins were quantified according to the protocol described previously (Rajput et al., 2022; Singh et al., 2024). Briefly, lyophilized samples (∼200 mg) were ground into powder and extracted with 80% methanol. The supernatant was filtered through a 0.22 µm PVDF membrane filter for further analysis. For acid hydrolysis, three volumes of 2 M acidic methanol: HCl were added to the supernatant and incubated at 90 °C for 45 min. The samples were dried in a rotavapor and resuspended in 1 mL of 80% methanol. The analysis was conducted on a 1290 Infinity II series UHPLC system (Agilent 290 Technologies, Santa Clara, CA, USA) equipped with a Zorbax Eclipse Plus C18 column kept at 30 °C. The appropriate parameters were as described previously (Singh et al., 2024). For the anthocyanin quantification, we used the LC–MS systems. The positive ionization voltage was set at 5500 V. Utilizing the Analyst software (version 1.5.2), the mass spectrometer was used in a variety of reaction monitoring modes for both qualitative and quantitative analysis. Analytical standards (Merck) were procured for each compound. Three biological replicates were used for anthocyanin quantifications.

### Dual-luciferase reporter assay

To test the transactivation potential of MtCBF4, the full-length CDS was cloned into the destination vector pBTdest (Baudry *et al.,* 2004) to generate CaMV *pro35S*-MtCBF4 constructs. Total genomic DNA of *M. truncatula* was isolated from A17 ecotype leaves using the DNeasy Plant Mini Kit (Qiagen). Promoters of *MtLAP1* (-2513 bp) and *MtJMJ13* (-2206) were amplified from genomic DNA of *M. truncatula* A17 ecotype and cloned into the p635nRRF vector (Kumar et al., 2018). For dual-luciferase reporter assays, reporter and effector constructs were transformed into *A. tumefaciens* strain GV3101 (pMP90) (GoldBio), and subsequently infiltrated into the abaxial side of *N. benthamiana* leaves according to the protocol described previously (Walter *et al.,* 2004). Protein extracts were prepared from infiltrated leaves 48 h post-infiltration, and LUC/REN values were measured using the Dual-Luciferase Reporter Assay System (Promega) according to the manufacturer’s protocol. Luminescence in the samples was quantified using the POLARstar Omega multimode plate reader (BMG Labtech). Values are given as means ± SD of the LUC/REN ratio of 3-4 independent biological replicates.

### Y1H assays

The Matchmaker Gold Yeast One-hybrid system (Clontech) and Y1H gold yeast strain were used. Promoter fragments having desired cis motifs were amplified and cloned into the pABAi vector (Takara Bio); Linearized pABAi plasmids carrying the promoter fragments were transformed into the Y1H Gold strain according to the Matchmaker protocol. MtCBF4 was sub-cloned into pGADT7-GW vector (Gal4-AD fusion), and were together with the empty vector into the Y1H strains previously transformed with AurR-reporter constructs (*proMtLAP1*-AurR, and *proMtJMJ13*-AurR). To check the interaction, the yeasts carrying both effector and reporter constructs were spotted onto medium containing a suitable concentration. of AbA per the manufacturer’s protocol. Growth of different yeast colonies was assayed using different concentrations of AbA to determine the minimum inhibitory concentration (MIC) values.

### ChIP-qPCR assay

ChIP assays were carried out as previously described with minor modifications (Saleh et al., 2008). Briefly, 1Lg of tissue each (EV and MtJMJ13-RNAi) samples were cross-linked with 1% formaldehyde to fix protein–DNA complexes. The samples were homogenized in liquid N2, followed by nuclei isolation and nuclei lysis. Sonication of chromatin samples was done on ice using a probe sonicator. Sonicated samples were first precleared with Protein A Agarose beads (Millipore; Burlington, MA, USA; #16-157), followed by incubating with H3K27me3 antibodies. The immuno-captured protein–DNA complexes were washed, and precipitated DNA samples were recovered.

RT-qPCR was conducted with 5 µL of 2x SYBR Green PCR Master mix (Applied Biosystems), 1 µL eluted DNA, and 0.5 µL of each forward and reverse primer, making a total volume of 10 µL. The relative fold enrichment of the different regions was calculated using the 2^−(–ΔCT)^ method (Livak & Schmittgen, 2001). The primer sequences used for ChIP-qPCR are listed in Supplemental Table S1.

### Electrophoretic mobility-shift assay

For the EMSA, MtCBF4 CDS was sub-cloned into the pGEX-4T-2 vector, and then transformed into *Escherichia coli* cells BL21 (DE3/Codon+) for protein induction. The culture was maintained up to a suitable OD_600_ of 0.5, and added with IPTG (final concentration 0.5 mM) and further cultured at 28 L for 12 h, followed by harvesting. GST-tagged recombinant protein was purified using Glutathione Resin (Catalog no. 786-310, G-Biosciences). The different probe sets containing intact and mutated DRE/CRT motifs were labelled with biotin using Pierce™ Biotin 3’ End DNA Labeling Kit (Catalog no. 89818, Thermo Fisher Scientific, USA). The EMSA was carried out using LightShift Chemiluminescent EMSA Kit (Catalog no. 20148, Thermo Fisher Scientific, USA) according to the instructions given in the kit manual. The primers and probe sequences for EMSA are shown in Supplementary Table S1.

### Plant protein isolation and Immunoblotting

For the in vivo demethylase activity assays, total proteins were extracted from *M. truncatula* hairy roots of *EV* and *MtJMJ13-RNAi* plants. Briefly, the root samples were ground and added to a cold protein isolation buffer. Supernatant was used for Western blot with suitable antibodies. The effect of *MtJMJ13* gene silencing on histone demethylation was estimated by immunoblotting with anti-H3K27me3 antibody. Rubisco was stained with Ponceau S for normalization.

EV and *MtCBF4-*overexpressing leaves were, likewise, used to isolate total proteins. The effect of MtCBF4 on histone methylation was determined by western blot with anti- H3K27me3 antibody. Rubisco was stained with Ponceau S for normalization.

### Statistical analysis

GraphPad Prism (version 9.0) was adopted to perform an un-paired t-test for all the comparisons. Asterisks indicate the level of significance: *P ≤ 0.05; **P ≤ 0.01; ***P ≤ 0.001, ****P ≤ 0.0001; ns, not significant.

## Acknowledgements

This work was supported by the core grant of the National Institute of Plant Genome Research and Department of Science and Technology-ANRF for Core Research Grant (CRG/2022/001178) to AP. NA acknowledges the Department of Biotechnology, Government of India, for Junior Research Fellowship. DC acknowledges support from the JC Bose fellowship (JCB/2020/000014) from Anusandhan National Research Foundation, Department of Science and Technology, Government of India. The authors are thankful to the DBT-eLibrary Consortium (DeLCON) for providing access to e-resources. We acknowledge the Metabolome facility at NIPGR for phytochemical analysis.

## Author’s contribution

JN and AP conceived the idea and designed the research. JN and NA conducted experiments. JN, NA, and AP interpreted the data. DC provides insightful suggestions during the discussion. JN wrote the manuscript. DC and AP improved the manuscript. AP coordinated the research project. All authors read and approved the final manuscript.

## Data availability

All data supporting the findings of this study are available within the paper and within the supplementary data published online.

## Declarations

The authors declare no conflict of interest.

---

**Figure S1. Transformation efficiency analysis in the infiltrated leaves of *M. truncatula*.** Semi-quantitative PCR for analyzing the expression of the *GFP* as a transformation marker in *EV* control, *MtCBF4-overexpressing,* and non-transformed leaves. *MtACTIN1* was used as a reference gene.

**Figure S2. Expression analysis of various biosynthesis genes in transiently overexpressing *MtCBF4* leaves vs control leaves.** Relative expressions of *MtPAL*, *MtCHS2, MtCHI*, *MtF3H*, *MtF3’H*, *MtF3’5’H*, *MtFLS1*, *MtIFS1*, *MtANR*, *MtLAR*, *MtGSTF7* and *MtWD40-1*, as determined by RT-qPCR. The data are mean values ±SD (n=3 biological), each derived from several hairy roots.

**Figure S3. Phylogenetic analysis of MtKFB1/2/3 of *M. truncatula* with previously characterized JMJ proteins from other species.** The phylogenetic tree was constructed in MEGA12 software using the maximum-likelihood method with 1000 bootstrap values and was visualized with iTOL v.6.4 (http://itol.embl.de/). Accessions used are MtKFB1 (Medtr8g069800.1), MtKFB2 (Medtr5g043750.1), MtKFB3 (Medtr5g033880), AtKFB01 (AT1G15670), AtKFB20 (AT1G80440), AtKFB39 (AT2G44130), AtKFB50 (AT3G59940), AtKFBCHS (At1g23390), CmKFB (MELO3C011980), VviKFB07 (Vitis01g00817).

**Figure S4. Transformation efficiency analysis in the hairy roots of *M. truncatula*.** RFP detection in the *EV* and *MtCBF4-silencing* hairy roots. Fluorescence as observed under fluorescence microscopy with RFP filter. Scale bar 500µm.

**Figure S5. Expression analysis of various biosynthesis genes, *MtCBF4* silencing *M. truncatula* A17 hairy roots vs control.** Relative expressions of *MtPAL*, *MtCHI*, *MtFLS1, MtF3H*, *MtF3’H*, *MtF3’5’H*, *MtFLS1*, *MtIFS1*, *MtANR*, *MtLAR*, *MtGSTF7*, *MtTT8* and *MtWD40-1*, as determined by RT-qPCR. The data are mean values ±SD (n=3 biological), each derived from several hairy roots.

**Figure S6. MSA of MtJMJ13 with its KMD4 orthologs.** Amino acid multiple sequence alignment of MtJMJ13 and selected KDM4 regulators from other plant species using MAFFT v.7.299b. The JmjN, JmjC, and C5HC2 domains are highlighted manually. Accessions used are MtJMJ13 (Medtr5g029370), AtJMJ13 (AT5G46910), SlJMJ3 (Solyc08g005240.3.1).

**Figure S7. Transformation efficiency analysis in the infiltrated leaves of *M. truncatula*.** Semi-quantitative PCR for analyzing the expression of the *GFP* as a transformation marker in EV control, *MtJMJ13-*overexpressing, and non-transformed leaves. *MtACTIN1* was used as a reference gene.

**Figure S8. Expression analysis of various biosynthesis genes in transiently overexpressing *MtJMJ13* leaves vs control leaves.** Relative expressions of *MtPAL*, *MtCHS2*, *MtCHI*, *MtF3H*, *MtF3’H*, *MtF3’5’H*, *MtFLS1*, *MtIFS1*, *MtANR*, *MtLAR*, *MtGSTF7* and *MtWD40-1* as determined by RT-qPCR. The data are mean values ±SD (n=3 biological), each derived from several hairy roots.

**Figure S9. Transformation efficiency analysis in the hairy roots of *M. truncatula*.** RFP detection in the *EV* and *MtJMJ13* silencing hairy roots. Fluorescence as observed under fluorescence microscopy with RFP filter. Scale bar 500µm.

**Figure S10. Expression analysis of various biosynthesis genes, *MtJMJ13* silencing *M. truncatula* A17 hairy roots vs control.** Relative expressions of *MtPAL*, *MtCHS2*, *MtCHI*, *MtF3H*, *MtF3’H*, *MtF3’5’H*, *MtFLS1*, *MtIFS1*, *MtANR*, *MtLAR*, *MtGSTF7*, *MtTT8*, and *MtWD40-1* as determined by RT-qPCR. The data are mean values ±SD (n=3 biological), each derived from several hairy roots.

**Figure S11. Differential accumulation of trimethylated histone H3K27 in *MtJMJ13*-silencing hairy roots compared to WT.** Immunoblotting of histone H3 with anti-H3K27me3 antibody in WT and *MtJMJ13*-silencing hairy roots. Ponceau S staining was used as a loading control.

**Figure S12. Expression analysis of various biosynthesis and regulatory genes in cold stress vs control *M. truncatula* plants.** Relative expression of anthocyanin biosynthesis genes like *MtPAL*, *MtCHS2*, *MtCHI*, *MtF3H*, *MtF3’H*, *MtF3’5’H*, *MtFLS1*, *MtIFS1*, *MtLAR*, *MtGSTF7, MtWD40-1, MtJMJ13, and MtKFB1/2/3* in the leaves of cold vs control plants at different time points. The values were normalized to *MtACTIN* expression. The data represent the mean ± SD of 3 biological replicates.

Supplementary File S1. Promoter analysis of the selected biosynthetic and regulatory genes Supplementary File S2. List of primers used in this study

