## Supplementary Figures for "Cold-responsive MtCBF4-MtJMJ13 positive feedback loop negatively regulates anthocyanin biosynthesis in *Medicago truncatula*"

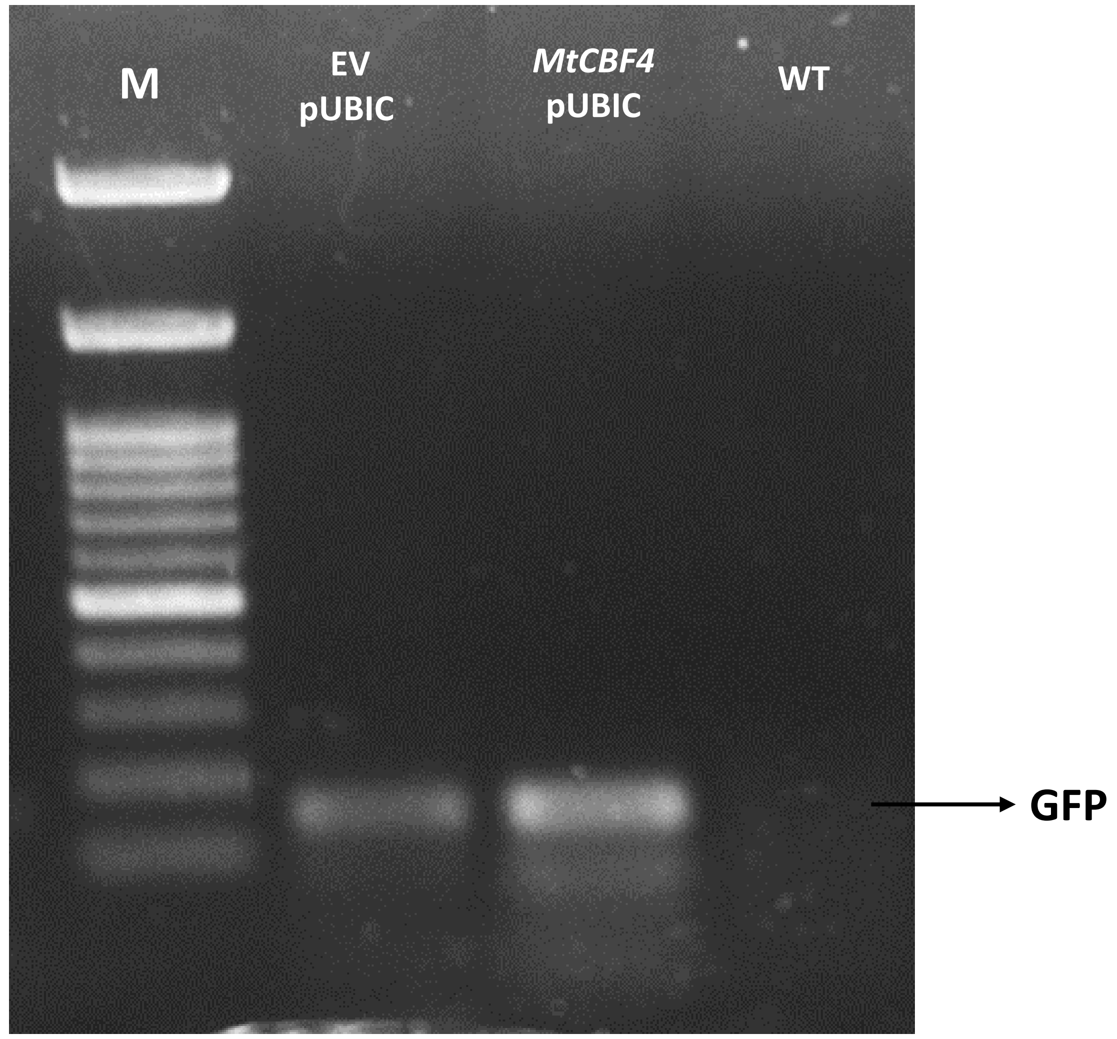
M
EV pUBIC
*MtCBF4* pUBIC
WT
GFP

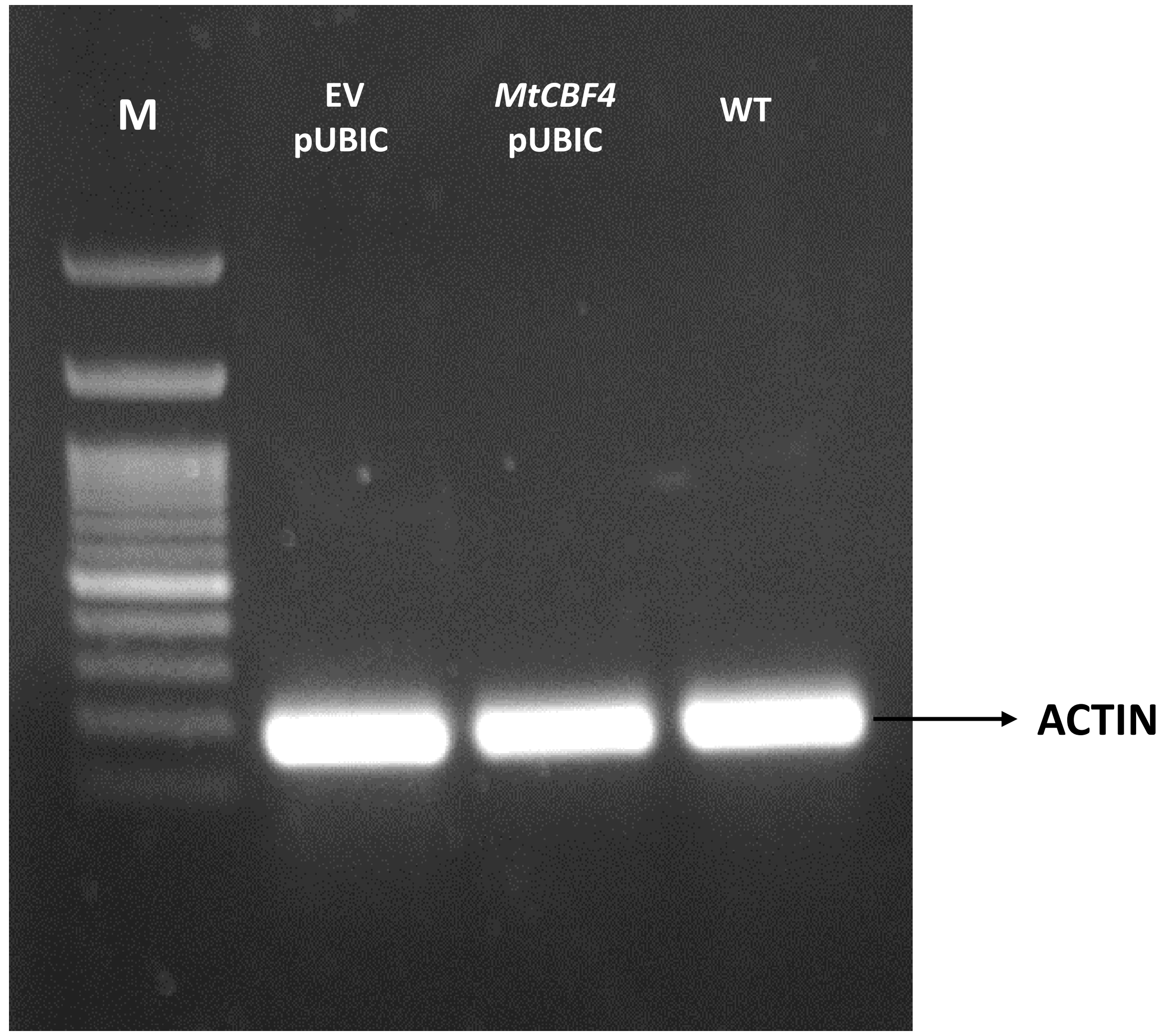
M
EV pUBIC
*MtCBF4* pUBIC
WT
ACTIN

**Fig. S1**

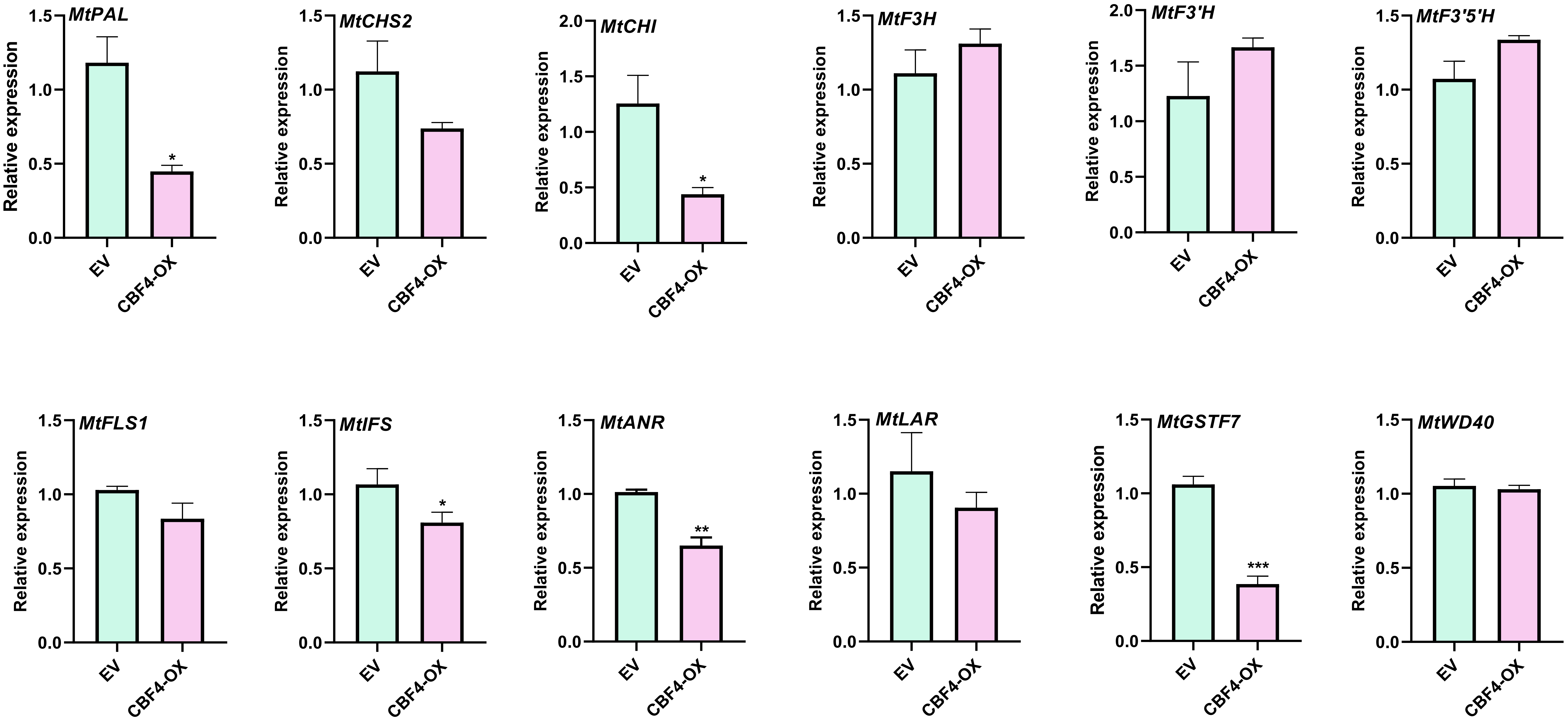
MtPAL
MtCHS2
MtCHI
MtF3H
MtF3'H
MtF3'5'H
MtFLS1
MtIFS
MtANR
MtLAR
MtGSTF7
MtWD40
Relative expression
EV
CBF4-OX
*
**
***
0.0
0.5
1.0
1.5
2.0

Fig. S2

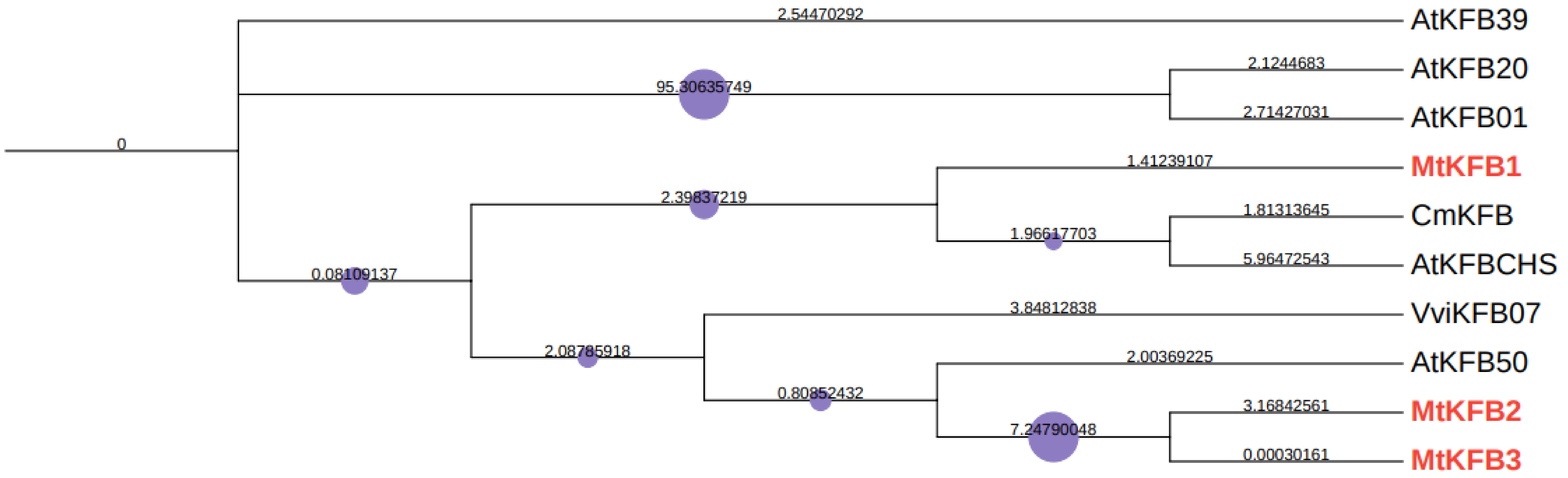

0
2.54470292
AtKFB39
95.30635749
2.1244683
AtKFB20
2.71427031
AtKFB01
1.41239107
MtKFB1
2.39837219
1.81313645
CmKFB
1.96617703
5.96472543
AtKFBCHS
0.08109137
3.84812838
VviKFB07
2.08785918
2.00369225
AtKFB50
0.80852432
3.16842561
MtKFB2
7.24790048
0.00030161
MtKFB3


**Fig. S3**

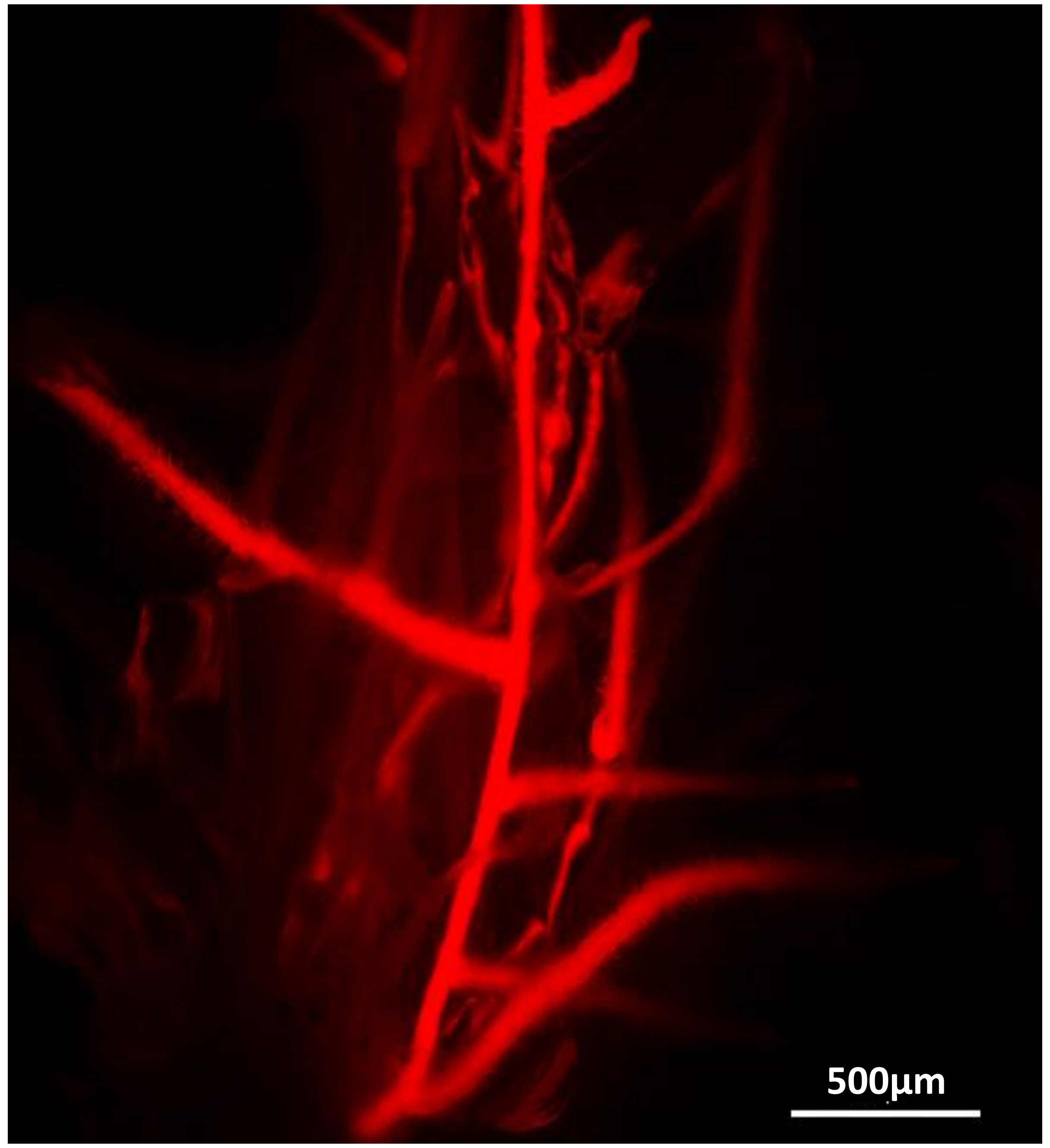


*EV-RNAi*

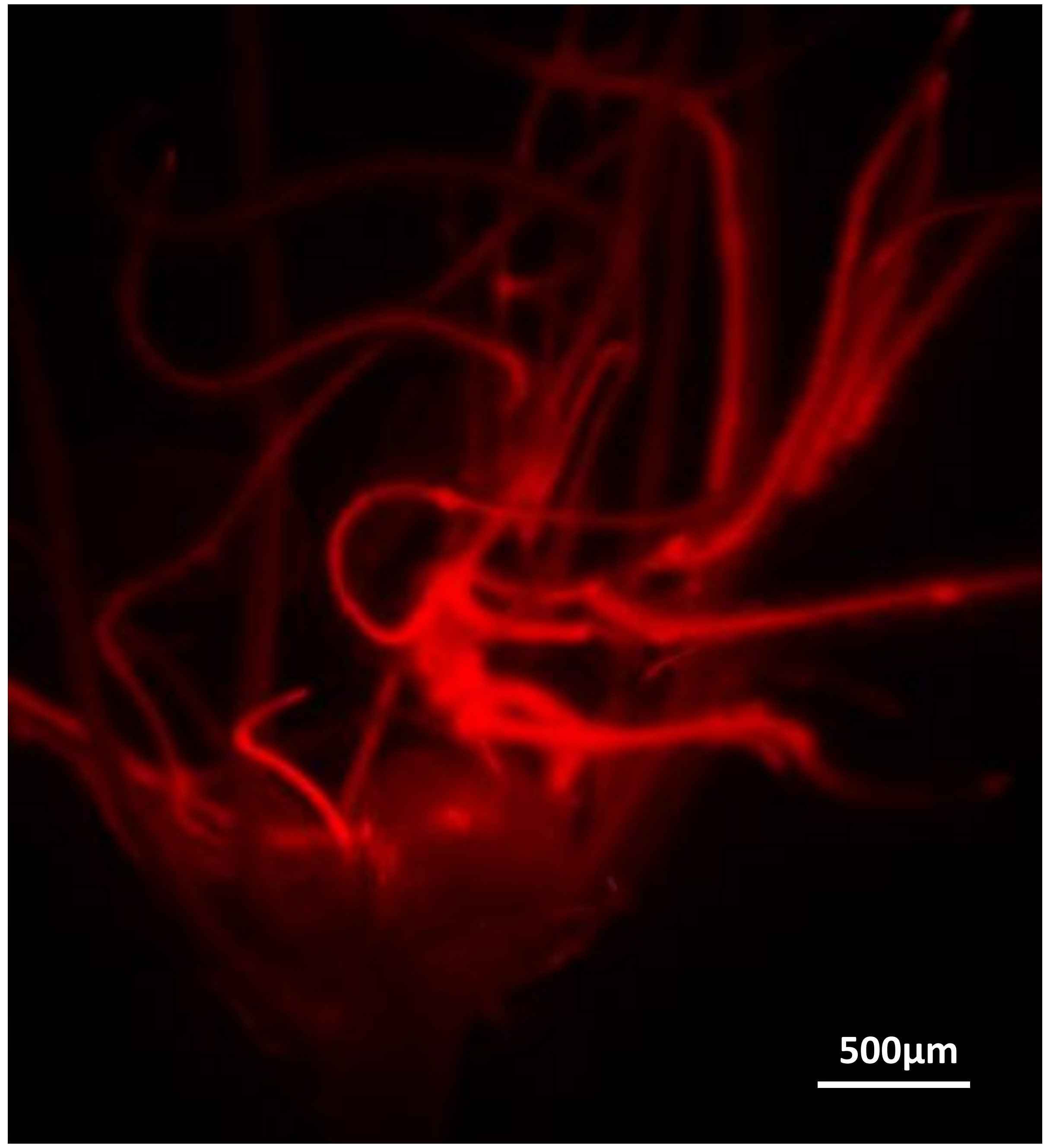


*MtCBF4-RNAi*

**Fig. S4**

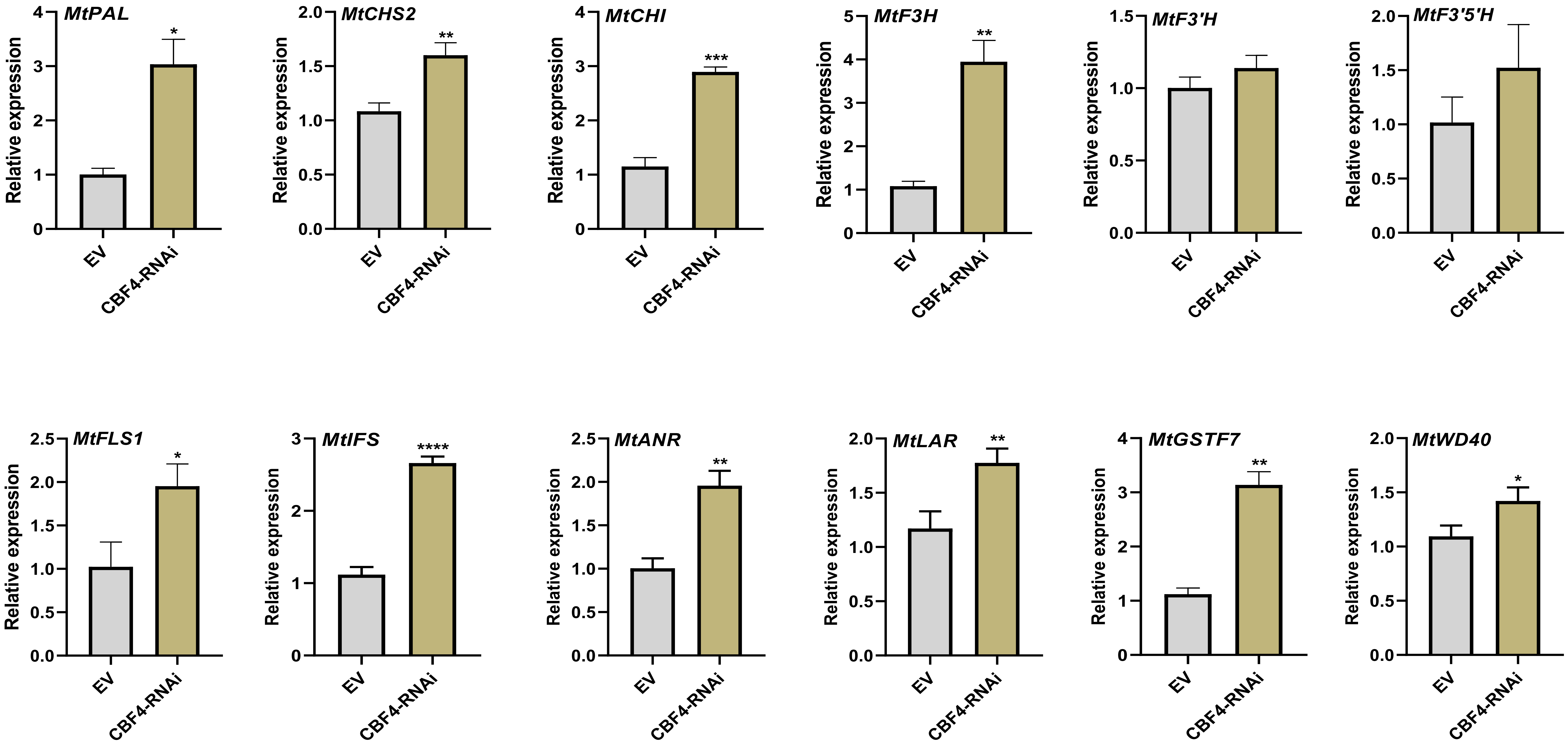

MtPAL
MtCHS2
MtCHI
MtF3H
MtF3'H
MtF3'5'H
MtFLS1
MtIFS
MtANR
MtLAR
MtGSTF7
MtWD40
Relative expression
EV
CBF4-RNAi
*
**
***
****


Fig. S5

```
AtJMJ13     -MAERRICLSKEAKDGLEFLKRKKLQKMRSDSVNETVGFSTMARSGGDALRPTSASCGMR
MtJMJ13     MVEGGRVRLSEEARNGLEFLKRKRLQRAQAIAATQTSAANVMNRSGGDALRG-SPSCGTR
SlJMJ3      ------------------------------------------MSRSGGDALRS-TASCGVR
                                                       * ********  : *** *

AtJMJ13     LRVTSSDTVSKVHGASTVRGGLMKEKVEKLETDDLKWTERLPECPVYRPTKEEFEDPLTY
MtJMJ13     LHGNP-------------DVFFKRKVDKFDTSDLDWTDKIPECPVYSPTKEEFEDPLVY
SlJMJ3      IRVNADMHS-GSGTSLNERNVFPKHKVAKFDTSDLEWTDKIPECPVYYPSKEEFEDPIVY
             :: .                .  : *.** *::*.**.**:::****** *:*******:.*
                    Jmj C
AtJMJ13     LQKIFPEASKYGICKIVSPLTATVPAGAVLMKEKSNFKFTTRVQPLRLAEWDSDDKVTFF
MtJMJ13     LQKIAPEASKYGICKIISPLSASVPAGVVLMKEQPGFKFTTRVQPLRFAEWDTEDKVTFF
SlJMJ3      LQKITPEASKYGICKIVSPIMASVPAGVVLMKEKVGFKFTTRVQPLRLAEWDRDDRVTFF
            **** ***********:**: *:*****.*****:  .***********:**** :*:****

AtJMJ13     MSGRTYTFRDYEKMANKVFARRYCSGGSLPDSFLEKEFWKEIACGKTETVEYACDVDGSA
MtJMJ13     MSGRNYTFREYEKMANKVFARRYCSVGCLPATYLEKEFWQEIGRGKMDTVEYACDVDGSA
SlJMJ3      MSGRNYTFRDFEKMANKVYARRYCSAGCLPPTYMEKEFWHEIASGKTESVEYACDVDGSA
            ****.****::*******:****** *.** :::*****:**. ** ::***********

AtJMJ13     FSSAPGDPLGSSKWNLNKVSRLPKSTLRLLETSIPGVTEPMLYIGMLFSMFAWHVEDHYL
MtJMJ13     FSTSPTDQLGNSKWNLKKLSRLPKSTLRLLETSIPGVTEPMLYIGMLFSMFAWHVEDHYL
SlJMJ3      FSSSPNDELGKCKWNMKRFSCLPKSVLRLLEKAIPGVTEPMLYIGMLFSMFAWHVEDHYL
            **::* * **..***::.* ****.*****.:****************************
                                         Jmj N
AtJMJ13     YSINYQHCGASKTWYGIPGSAALKFEKVVKECVYNDDILSTNGEDGAFDVLLGKTTIFPP
MtJMJ13     YSINYQHCGASKTWYGIPGHAALEFERVVREHVYSTDILSSDGEDGAFDVLLGKTTLFPP
SlJMJ3      YSINYHHCGAAKTWYGIPGHAALDFEKVVRENVYNNDILTADGEDGAFDVLLQKTTFFPP
            *****:****:********* ***.**:**:* **. ***::*********** ***:***

AtJMJ13     KTLLDHNVPVYKAVQKPGEFVVTFPRAYHAGFSHGFNCGEAVNFAMGDWFPFGAIASCRY
MtJMJ13     NILMEHKVPVYKAVQKPGEFVITFPRAYHAGFSHGFNCGEAVNFALGDWFPLGAIASRRY
SlJMJ3      NILSEHDVPVYKAVQKPGEFIVTFPRAYHAGFSHGFNCGEAVNFATGDWFPIGSIASRRY
             : * :*.**************::*********************** *****:*:*** **

AtJMJ13     AHLNRVPLLPHEELICKEAMLLNSSSKSENLDLTPTELSGQRSIKTAFVHLIRFLHLARW
MtJMJ13     ALLNRVPLLPHEELLCKEAMLIHSSLELEDSDFPSSDLLSHHRTKISFINLLRFQHCASW
SlJMJ3      ALLNRVPLLPNEELLCKEAMLLLTDLELEYSAISSADLITHHTIKVSFINLMRFHHRARW
            * ********:***:******:  :. : *   :  ::*   ::  * :*::*:** * * *
                                                        ZnF
AtJMJ13     SLMKSGLCTGLVSNTYGTIVCSLCKRDCYLAFINCECYSHPVCLRHDVKKLDLPCGTTHT
MtJMJ13     LLMKSRACISVSSHSHGTILCSLCKRDCYVAYVDCSCHMHPVCLRHDVKSLDFICGSKHT
SlJMJ3      CFLKLKAFSGISSFSHSTILCSICKRDSYVAYLNCSCYSHAACLRHDPRSLHFPCGSSRT
              ::*     .: * ::.**:**:****.*:*::*.*: * .***** :.*.: **:.:*

AtJMJ13     LYLRDNIEDMEAAAMKFEKEDGVSDLIT----TDEDLYKYPSSITLPAAKEDGYTPYSTI
MtJMJ13     LYLREDIADMEAAAKMFEQEDGILDEISKQSKSDQNMYSHPLSDMFQRAEANGYEPYCEL
SlJMJ3      LCLREDILDIETTARKFELDDNVLHDVAHYQEGDDL---ALLLNMFPQAEEEGYVPYCEI
            * **::* *:*::*  ** :*.: . ::    *:           :  *: :** **. :
```

Fig. S6

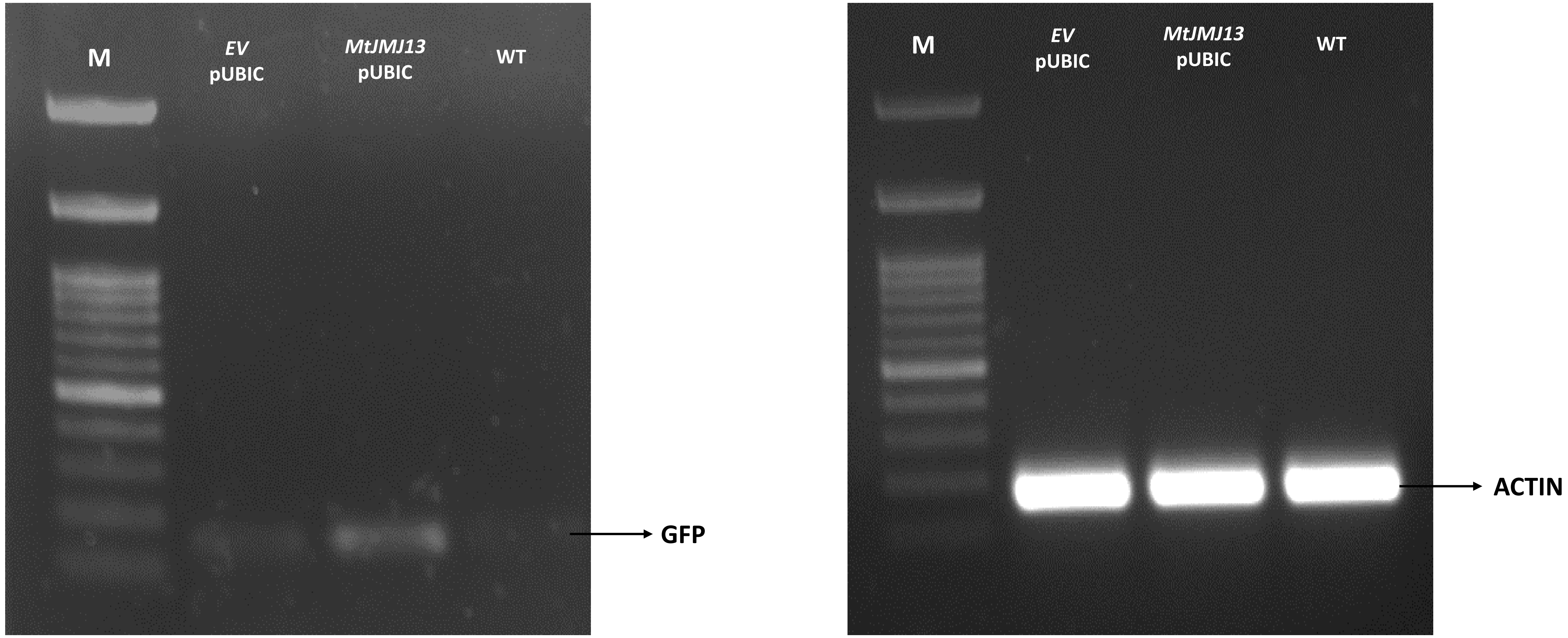
M
*EV* pUBIC
*MtJMJ13* pUBIC
WT
GFP
M
*EV* pUBIC
*MtJMJ13* pUBIC
WT
ACTIN

Fig. S7

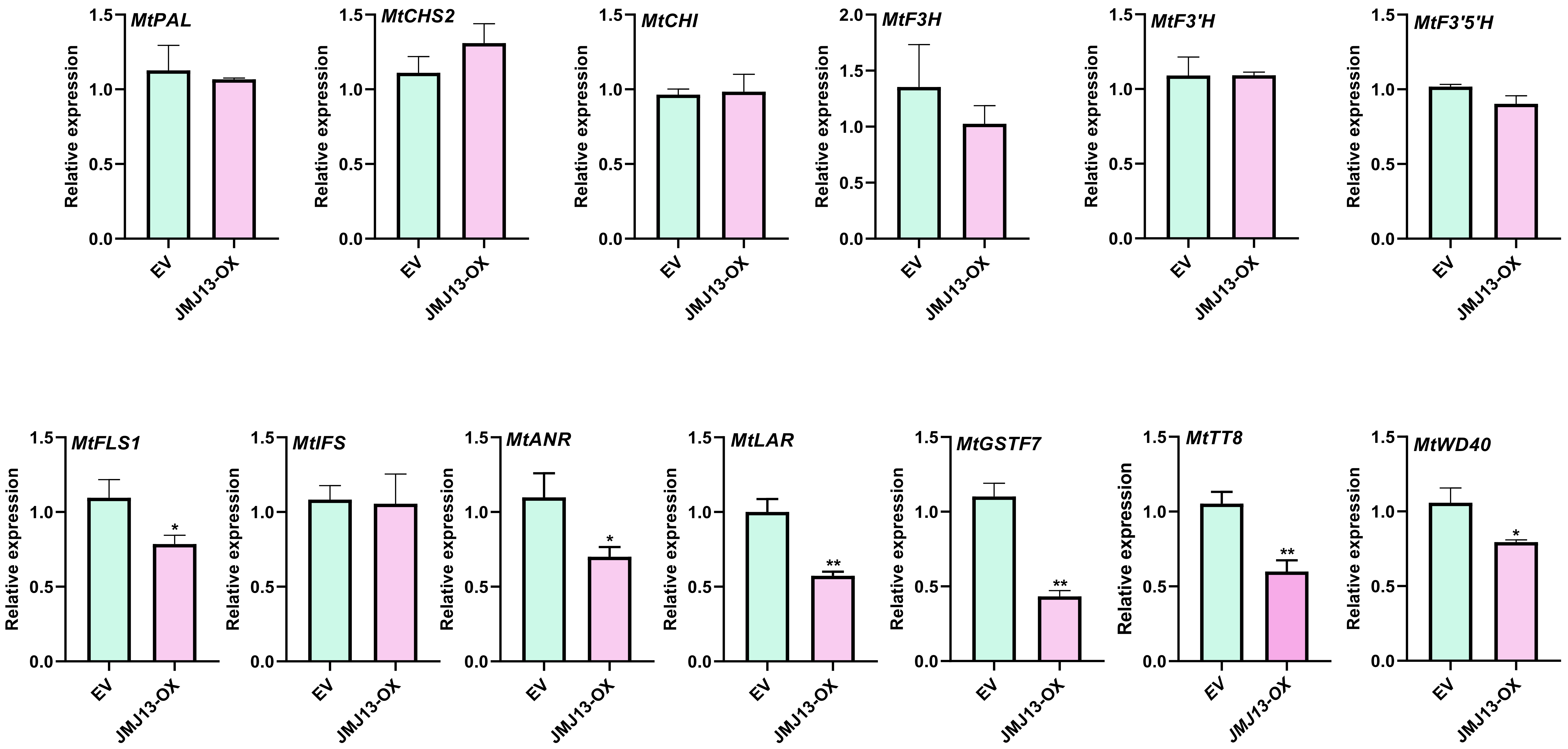

MtPAL
MtCHS2
MtCHI
MtF3H
MtF3'H
MtF3'5'H
MtFLS1
MtIFS
MtANR
MtLAR
MtGSTF7
MtTT8
MtWD40
Relative expression
EV
JMJ13-OX
*
**


Fig. S8

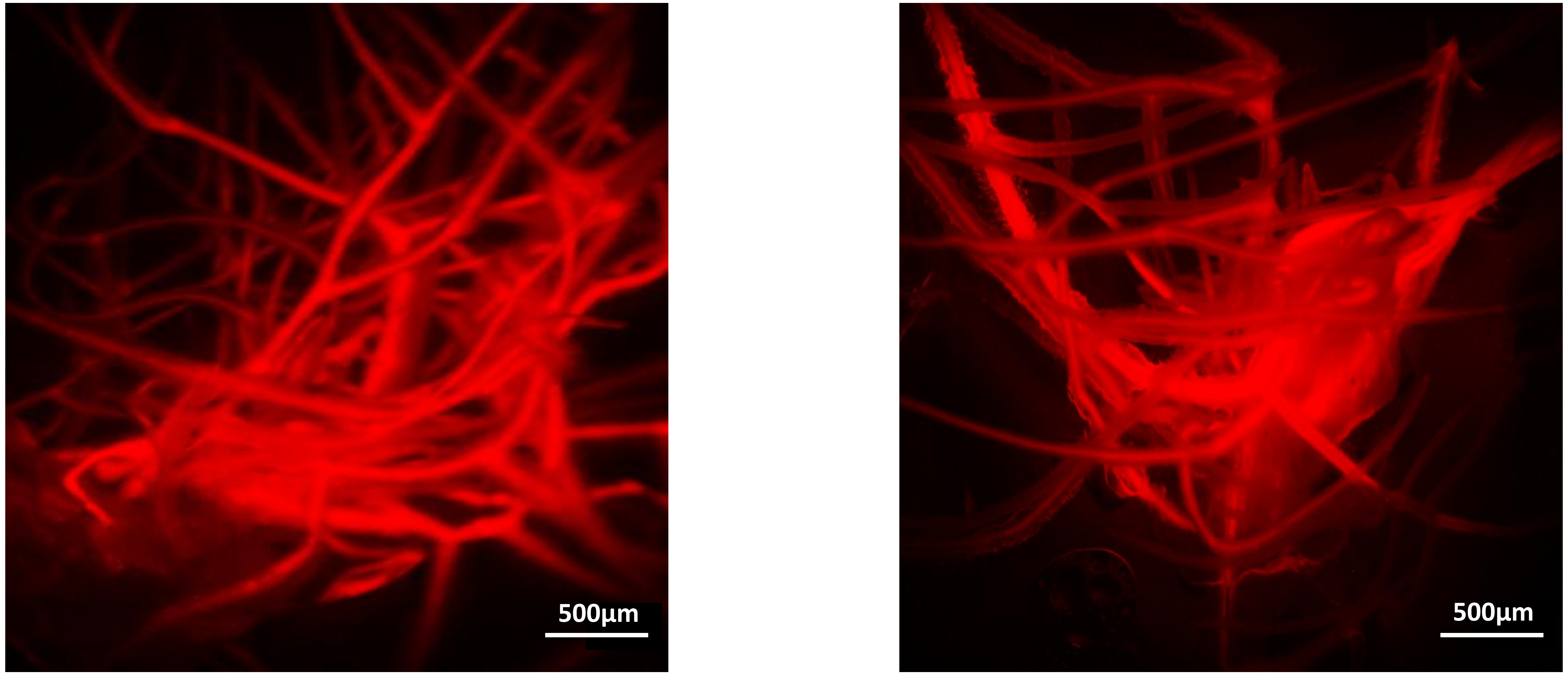


*EV-RNAi* *MtJMJ13-RNAi*

**Fig. S9**

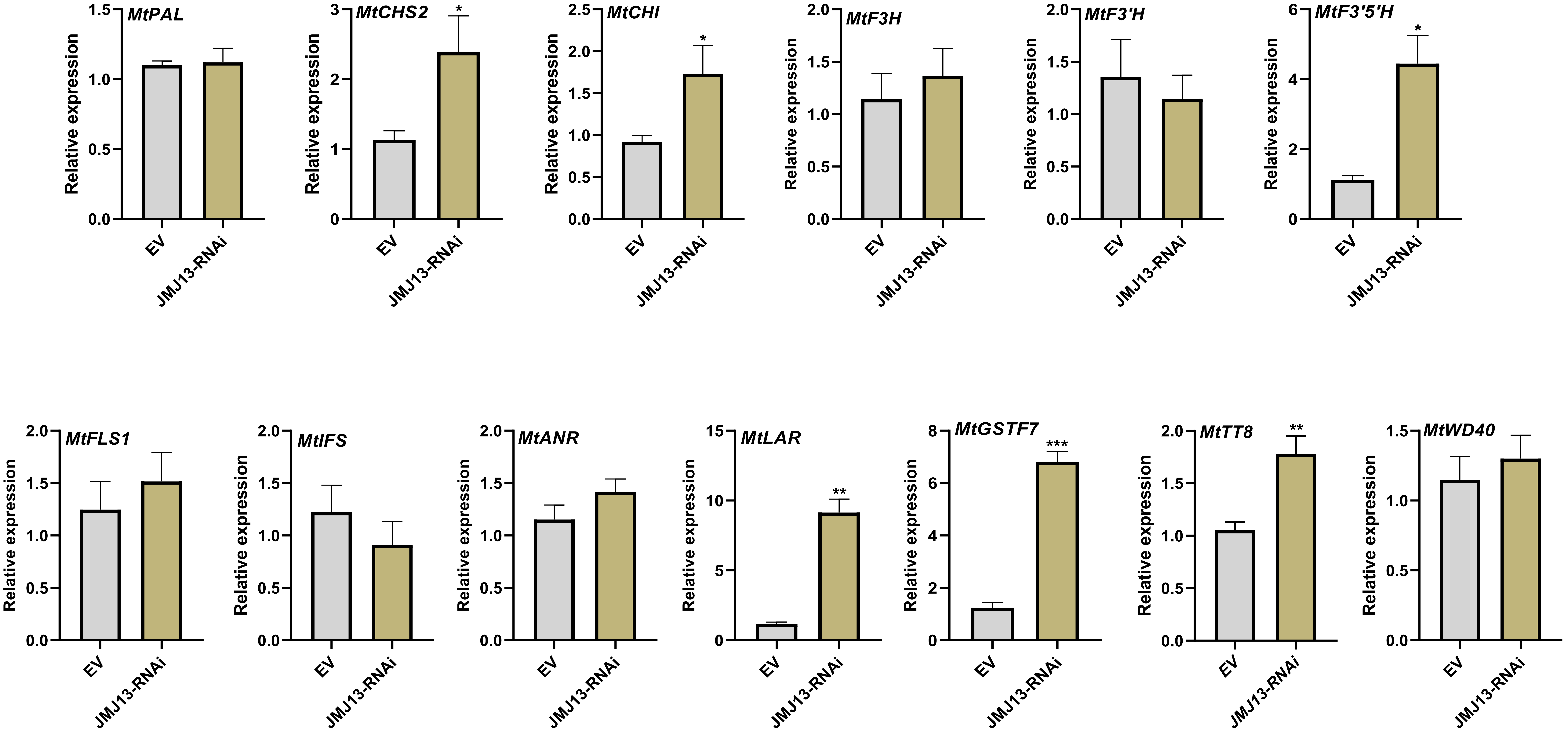
MtPAL
MtCHS2
MtCHI
MtF3H
MtF3'H
MtF3'5'H
MtFLS1
MtIFS
MtANR
MtLAR
MtGSTF7
MtTT8
MtWD40
Relative expression
EV
JMJ13-RNAi
*
**
***

Fig. S10

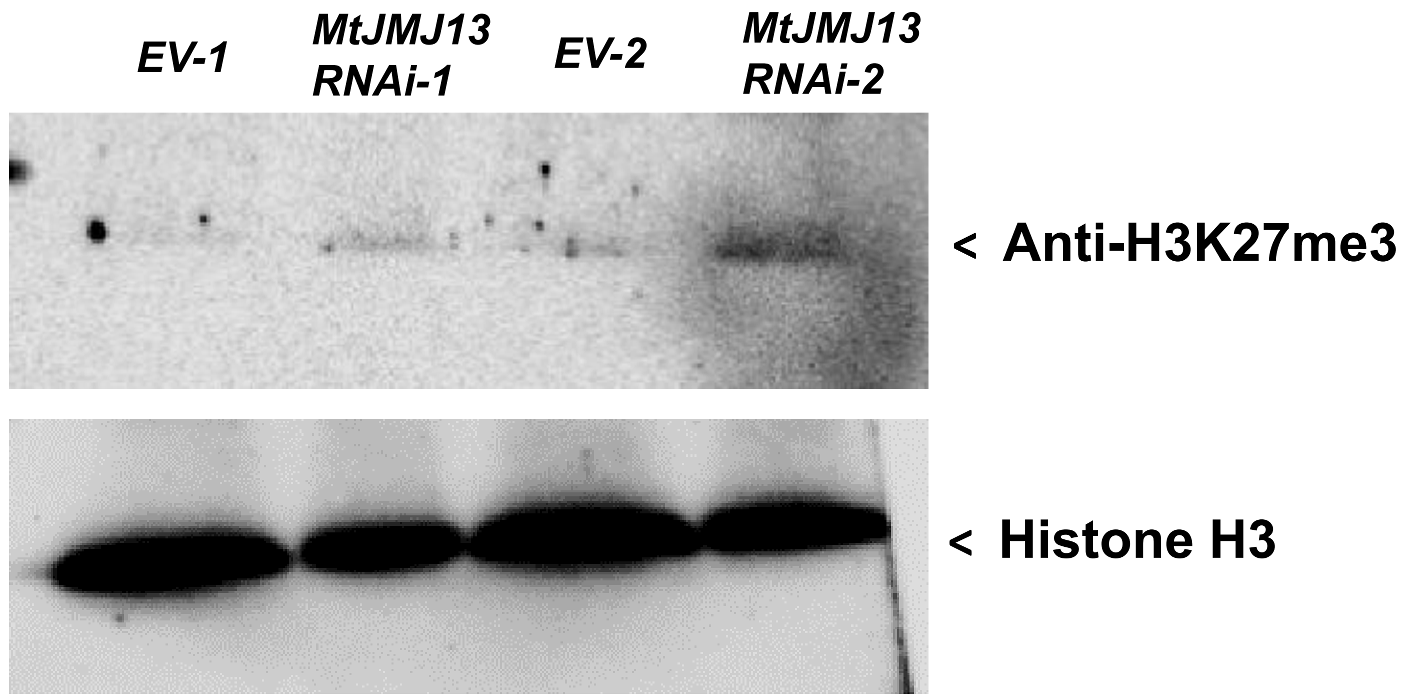
EV-1
MtJMJ13 RNAi-1
EV-2
MtJMJ13 RNAi-2
< Anti-H3K27me3
< Histone H3

Fig. S11

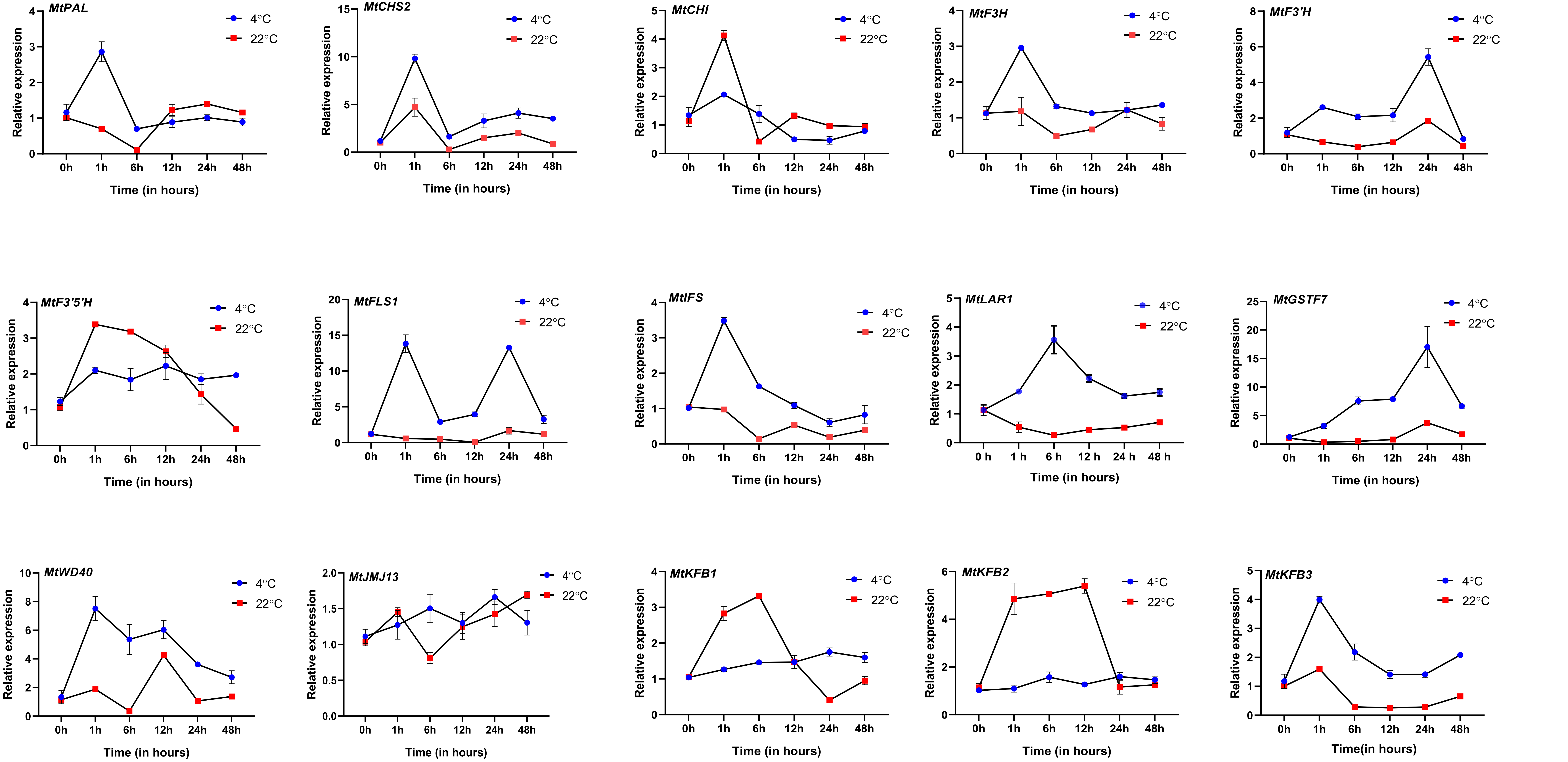
MtPAL
MtCHS2
MtCHI
MtF3H
MtF3'H
MtF3'5'H
MtFLS1
MtIFS
MtLAR1
MtGSTF7
MtWD40
MtJMJ13
MtKFB1
MtKFB2
MtKFB3
4°C
22°C
Relative expression
Time (in hours)
0h
1h
6h
12h
24h
48h
0 h
1 h
6 h
12 h
24 h
48 h
Time(in hours)

Fig. S12
