## Supplementary Table S1 for "Cold-responsive MtCBF4-MtJMJ13 positive feedback loop negatively regulates anthocyanin biosynthesis in *Medicago truncatula*"

| **Supplementary Table S1: List of primers used in the present study** |
| --- |

| **Primer Name** | **Sequence (5’ to 3’)** | **Purpose** |
| --- | --- | --- |
| MtJMJ13_For | CACCATGGTAGAAGGAGGTAGAG | Forward primer for cloning of MtJMJ13 in pENTR-dTOPO vector |
| MtJMJ13_Rev | ATTCAATCTGCTCTCTATGC | Reverse primer for cloning of MtJMJ13 in pENTR-dTOPO vector |
| MtCBF4_For | CACCATGAATCAATTTTCGGAGTCAC | Forward primer for cloning of MtCBF4 in pENTR-dTOPO vector |
| MtCBF4_Rev | AAATGAGTAACTCCACAATG | Reverse primer for cloning of MtCBF4 in pENTR-dTOPO vector |
| MtCBF4_For | TAGGATCCATGAATGAATCAATTTTCGGAGTC | Forward primer for cloning of MtCBF4 in protein expression vector (BamHI) |
| MtCBF4_Rev | ATCTCGAGTTAAAATG AGTAACTCCACAATG | Reverse primer for cloning of MtCBF4 in protein expression vector (XhoI) |
| proMtJMJ13_For | CACCATCAAGATGGGACACCCTTG | Forward primer for cloning of MtJMJ13 promoter for Dual luciferase assay |
| proMtJMJ13_Rev | TTCATTACGAACTCGTTCCC | Reverse primer for cloning of MtJMJ13 promoter for Dual luciferase assay |
| proMtLAP1_For | CACCCAGATGACCATCTATGTAAGTC | Forward primer for cloning of MtLAP1 promoter for Dual luciferase assay |
| proMtLAP1_Rev | ATATAGATATATGCTCCTTTGC | Reverse primer for cloning of MtLAP1 promoter for Dual luciferase assay |
| MtJMJ13RNAi_For | CACCCAAATATCCAACAGTGCATGC | Forward primer for cloning of MtJMJ13 for RNAi |
| MtJMJ13RNAi_Rev | ATTCAATCTGCTCTCTATGC | Reverse primer for cloning of MtJMJ13 for RNAi |
| MtCBF4RNAi_For | CACCAACGGGAGGAGGAAGAG | Forward primer for cloning of MtCBF4 for RNAi |
| MtCBF4RNAi_Rev | CTAGTACGAACTTCCTGCAC | Reverse primer for cloning of MtCBF4 for RNAi |
| MtCHS2_For | CGAAGAATGGGGTCAACCGAAGT | RT-qPCR forward primer |
| MtCHS2_Rev | CAAACGAAGTACTGTTCCACCTGC | RT-qPCR reverse primer |
| MtCHI1_For | GGCGACAGCAGCACCCACAAT | RT-qPCR forward primer |
| MtCHI1_Rev | CTCCATCAATATCTAACCCCCTCAC | RT-qPCR reverse primer |
| MtJMJ13_For | AGTCCTTCCTGAAGGAAAAAG | RT-qPCR forward primer |
| MtJMJ13_Rev | CCCTTGCAAATCTGTTCTTC | RT-qPCR reverse primer |
| MtCBF4_For | CGGAAGCAAGGGATATTCAA | RT-qPCR forward primer |
| MtCBF4_Rev | CAAATTCCACGTCAGCAACA | RT-qPCR reverse primer |
| MtKFB1_For | TAGGTAACGCAGAAAACATG | RT-qPCR forward primer |
| MtKFB1_Rev | CGTATTCAACAATTCACTTTTG | RT-qPCR reverse primer |
| MtKFB2_For | CAATCAATGCCGAAAAAGATG | RT-qPCR forward primer |
| MtKFB2_Rev | CTTCCCATTTCATTGCAATAGC | RT-qPCR reverse primer |
| MtKFB3_For | CTGCTTTACACCTTATCCCC | RT-qPCR forward primer |
| MtKFB3_Rev | CAACGCGCGCTACTATTG | RT-qPCR reverse primer |
| MtMYB2_For | TAATCACAAACTACACAAAGGC | RT-qPCR forward primer |
| MtMYB2_Rev | CACACGACTTGATGGAGAG | RT-qPCR reverse primer |
| MtTT8_For | CACCTGAGGACCTAACAGAATC | RT-qPCR forward primer |
| MtTT8_Rev | GCTATCCACTTCATTTGCACC | RT-qPCR reverse primer |
| MtF3H_For | GGAGAGAAATCTGTGATGGAAG | RT-qPCR forward primer |
| MtF3H_Rev | AAGTCCCTAAGTTCTTTTTCTTCC | RT-qPCR reverse primer |
| MtF3’H_For | GGAACTTGAAAATGGACTCAATG | RT-qPCR forward primer |
| MtF3’H_Rev | AGAAACAAGACGAGTACACGTG | RT-qPCR reverse primer |
| MtF3’5’H_For | AGACATGGTTTTCGCGGACT | RT-qPCR forward primer |
| MtF3’5’H_Rev | CCCATTTCGTCCCCTCGAAT | RT-qPCR reverse primer |
| MtIFS_For | CAATCCTCCGAGTCCCAAAC | RT-qPCR forward primer |
| MtIFS_Rev | GGGTTATCCAAAAGGTGAAGATGA | RT-qPCR reverse primer |
| MtFLS_For | CAATCCAAAGATGCTACATCAACAA | RT-qPCR forward primer |
| MtFLS_Rev | ATAGGTACTTCGAGTTTGACACC | RT-qPCR reverse primer |
| MtActin_For | ACTCACACCGTCACCAGAATCC | RT-qPCR forward primer |
| MtActin_Rev | TCAATGTGCCTGCCATGTATGT | RT-qPCR reverse primer |
| MtANR_For | CAACTTCTGGTCGATACATTTGC | RT-qPCR forward primer |
| MtANR_Rev | CTGAGGGTATCGTTTGCTGAG | RT-qPCR reverse primer |
| MtLAP1_For | CCCCAACATCAACAGAGGAAGA | RT-qPCR forward primer |
| MtLAP1_Rev | TTCTTTGCCAAATTTGTGTGCC | RT-qPCR reverse primer |
| MtANS_For | GGTTGGAAGGTGGAAGGTTA | RT-qPCR forward primer |
| MtANS_Rev | CCCATTTGCCCTCATAGAAA | RT-qPCR reverse primer |
| MtDFR1_For | TCGTCCACTTGGATGATCTTTG | RT-qPCR forward primer |
| MtDFR1_Rev | CTCCCTTCTACTTCCATATGCTCAA | RT-qPCR reverse primer |
| MtPAL_For | TTGCCAAAAGAGGTTGAAAGTG | RT-qPCR forward primer |
| MtPAL_Rev | TGTTTGGAATTGTTGGGTTTCC | RT-qPCR reverse primer |
| MtLAR_For | TTTCCACAGCACATCCAACCTA | RT-qPCR forward primer |
| MtLAR_Rev | GACAATGGCACCTTTCTCTTGG | RT-qPCR reverse primer |
| MtGSTF7_For | CTCCAGCCCTTTGGTCAAGTT | RT-qPCR forward primer |
| MtGSTF7_Rev | ACGGTCTGCATACTTTGTTGC | RT-qPCR reverse primer |
| proMtLAP1_F1F | AGATGAGTCTCCCGACACTACGGTAGCCAC | Forward probe of the F1 site for the EMSA assay |
| proMtLAP1_F1R | GTGGCTACCGTAGTGTCGGGAGACTCATCT | Reverse probe of the F1 site for the EMSA assay |
| proMtLAP1_F2F | GTCTCTATTCTTTACCGACTCAAATCTAACA | Forward probe of the F2 site for the EMSA assay |
| proMtLAP1_F2R | TGTTAGATTTGAGTCGGTAAAGAATAGAGAC | Reverse probe of the F2 site for the EMSA assay |
| proMtJMJ13_F1F | ATCCTGTACAAGTCGGCAGTTCAACAGAAA | Forward probe of the F1 site for the EMSA assay |
| proMtJMJ13_F1R | TTTCTGTTGAACTGCCGACTTGTACAGGAT | Reverse probe of the F1 site for the EMSA assay |
| proMtJMJ13_F2F | TTTCAATGATGTCGATTTTTTTGTTG | Forward probe of the F2 site for the EMSA assay |
| proMtJMJ13_F2R | CAACAAAAAAATCGACATCATTGAAA | Reverse probe of the F2 site for the EMSA assay |
| MtCBF4F1_For | CACAAACTTTCTGACAAAAC | ChIP-qPCR forward primer |
| MtCBF4F1_Rev | GTGTTCTGTCAATGAGGAG | ChIP-qPCR reverse primer |
| MtCBF4F2_For | CCACACTGTTTTTCAACTAG | ChIP-qPCR forward primer |
| MtCBF4F2_Rev | CTCTGCAAGAAACCTCTCTG | ChIP-qPCR reverse primer |
| MtCBF4F3_For | GTCGCTGCGATTGCACTG | ChIP-qPCR forward primer |
| MtCBF4F3_Rev | CTCCGGCCTGAAAGCCTC | ChIP-qPCR reverse primer |
| MtKFB2F1_For | TCTATAAACATGTGGGCAGC | ChIP-qPCR forward primer |
| MtKFB2F1_Rev | AATGGCTGCATCTTAAGGG | ChIP-qPCR reverse primer |
| MtKFB2F2_For | CCGTGAAGCCGTGGCTC | ChIP-qPCR forward primer |
| MtKFB2F2_Rev | ACCGTAAAGCTGGGGTGTG | ChIP-qPCR reverse primer |
| MtKFB2F3_For | GATACCACAGCGTCGATG | ChIP-qPCR forward primer |
| MtKFB2F3_Rev | CGCTCTCTTCTCCTCGTAG | ChIP-qPCR reverse primer |
| MtKFB2F4_For | ACCAGAAGAGATGGTGGC | ChIP-qPCR forward primer |
| MtKFB2F4_Rev | CATCACCGGCGCACAATAC | ChIP-qPCR reverse primer |
| MtKFB3F1_For | CACACCCGAAACTCCATTAC | ChIP-qPCR forward primer |
| MtKFB3F1_Rev | CATGAGGGAGAATATAGCTTC | ChIP-qPCR reverse primer |
| MtKFB3F2_For | ACCAAAGCTAGTGAATTCCC | ChIP-qPCR forward primer |
| MtKFB3F2_Rev | TGCATGGTGCCATTTAAGATG | ChIP-qPCR reverse primer |
| MtKFB3F3_For | TTGTGACGGAGAAAGATTCCG | ChIP-qPCR forward primer |
| MtKFB3F3_Rev | CCATCAACTTCTCCGATGAAG | ChIP-qPCR reverse primer |
| MtKFB3F4_For | TTCAGAGTTTGGTTCAGTTG | ChIP-qPCR forward primer |
| MtKFB3F4_Rev | CCTCACACTCCTCCACTC | ChIP-qPCR reverse primer |
| MtMYB2F1_For | CATCAACAAAGGTGCTTG | ChIP-qPCR forward primer |
| MtMYB2F1_Rev | ATATAGTATGAGTGAATGAGATG | ChIP-qPCR reverse primer |
| MtMYB2F2_For | GAAGACCTCATCATCAAACTC | ChIP-qPCR forward primer |
| MtMYB2F2_Rev | GAAACGTGCTTTTAGACAC | ChIP-qPCR reverse primer |
| MtMYB2F3_For | GATTTTCTCAAAGAATTCGTC | ChIP-qPCR forward primer |
| MtMYB2F3_Rev | CCCTAAAATTAACTAACTGTAC | ChIP-qPCR reverse primer |
| MtMYB2F4_For | GGTTGCCTGGACGTAC | ChIP-qPCR forward primer |
| MtMYB2F4_Rev | GTAGGAAAGCCTTTGTGTAG | ChIP-qPCR reverse primer |
| proMtJMJ13_For | GGATCCATGGTAGAAGGAGGTAGAGTAC | Forward primer for cloning of MtJMJ13 promoter in pAbAi vector for Y1H assay (BamHI) |
| proMtJMJ13_Rev | CTCGAGAATAGCCCCAAGAGGAAACC | Reverse primer for cloning of MtJMJ13 promoter in pAbAi vector for Y1H assay (XhoI) |
| proMtLAP1_For | AAGCTTCTCCACCCTTTGTTGAAGAG | Forward primer for cloning of MtLAP1 promoter in pAbAi vector for Y1H assay (HindIII) |
| proMtLAP1_Rev | CTCGAGCATTGCTCGACATCTGAATTTG | Reverse primer for cloning of MtLAP1 promoter in pAbAi vector for Y1H assay (XhoI) |
