## Supplementary File S1 for "Cold-responsive MtCBF4-MtJMJ13 positive feedback loop negatively regulates anthocyanin biosynthesis in *Medicago truncatula*"

*MtJMJ13pro*

ATCAAGATGGGACACCCTTGAAAACCCTATTGTAAAGTTACAAAGATACAATTTTCGATAATAAGAACTTATGATTTAACATAAAAATTTATTTATTTAGATAAATTATTTTTTATAAGTTAAAAAATAATCCAATTCAAATTGGTCATAAATAAATAAAATTACTTGAATAATTTCATATTTATGTGTCGTGTTATGAGAAATGTTATATGTACAAACATTTTGTAAAAAAAAAATAAAATTATTTTAGTACGATTTTCTCTCTCATCTTCTCATTATCTTTTTATTTTCTCTTTTATTTTTCTTTGTTGTTTTTGCTCAAATAAAAGGAAGAACAAAAAAGTTGAATAAACAATTATCACAAAATCGTTGTACAAACATCAATTCTCAAATACTTTCGATCAAAAACATCGTTGCTACAGATATGTCGCGTCCGATGACTATGTTAGGTTAGGAGTAAGTACCTCATAGCACTTACAATACGGAAAAGTCACTGTGCATGTGACAACCCCGTTGAAAAATCCATTAATACCCTTTATTCTCGCGCCAAGATGAACCAACTGAACAAGGACAACCTCGTCCTTTTCTTTCTACAGTCTTAACTAACTGGAAGGTAGGTGAGAGTTATCCTGTACAA**GTCGG**CAGTTCAACAGAAAAAAGTACACATGCCTCTGAGCCTTTCTGATTCGAGGGAGTTTTCAGTTCAAGTACAATTCAACGGTTCAAAGTAAACGACACAAAATATCAACGGTTAGATGTCAAACTTCATCAAAGTCATATATATATTCCACCAAAGTCTTATAACACGTGATTATTCTTTTTTTCTGCCACTTTATTTTCCTCTGACCGATACAGAAGCAGAACCTCATCGATAACACCGCTTTTTTCTTTTACCAAAATACCCTCCACAACATGACATTAATTACACGCTCTTTAATTCTCTACCCGCCAATAAATACTCCTAACAGTAACGGTGATTGTGTTTCCGTCCTATTCGCGTAACGGCGTTATATTTAATCCGTGTTTACACTCATAACGCGTTTTGCATTTTGGACGGCGTGTTTGTTTCGTTGAACCACCTAACAACCACCACTTCTCCTCAAAGTTTCTTTTGATTTAATTATTTTATTCAACTAAACAAACCAAAAAATAAATAAACGAACTAAAAGGTACCTTAAAAATGAAGAAGAAATTAAAGTATATTTGATTTGAGTGTAAAAAAAATGAAAATAAAAGTTGTTAATAAATTCTAATCAATATTACTAGGTCAAATCTATTCACATGAATTCTTCAAATTCAAACTTTAGACTACACAGTGGATATTGTTAGATTAAACACAATCATTCTATTTTAAATTTGATATTTGTGTCAAAATTTCAATGAT**GTCGA**TTTTTTTGTTGTACACATTAATTCTTACTCAATATATAAAATGAATTAATTTGTGCTTGATATAAAAATGTTAAGGTTTCTTCAATTTTATGTCTATAAAGATTAAATTGACTCGCGGAAAAATAAAATAAAATAATAAAATTGATAAATATTATTTATTGAGTTTGTTGCATTTGAAAGTCATCACAATCTATATGACTCTTTTTAGTGTAAAATTTATATGATTTTTTTGGTGGTGTAAATCTATATGATTTGTTACACGCGGAAGCTACTGAAAAAAAAAAAAAAATTCCAATATATATTCGGGTTTTCAGAGCAGAAAAAGCGATTAATTAATTTGGAGTATAAAACTATAAATTATCACATCGTTACCCTTTTTACGTTTTCTAAGCATTCATATCTTTTCCTTCTCTCTCCTCATCACTGTTTCTCTCCGTTTTCTTTTCTCGCCGGGAAACAAACTTTTCCGTTCACCGGAAAACCGCTTTCTCTTTTTTTCTTCTATATTCCGCGCCACCGTAGGAAGCCGGTCACGGTGTTCCGATTGAAACACTCGCATTCACTCTCAATTCCCCGTGTTTGCTCCTACACGGCGTTGGATCTCTCCGTAACGCCGTATCACCCTCGTATACGGCGTCGTTCGAGAATGTGGCGAGCCTCGTTGTATTAAAACGCTACGTTCTAATCTACAACGTTCAGGATCGTGATTTAGGGTTTCGCTTTGTGATTTCGTTTCGGTTTCGTGATTCTTCAAACTTGTGATTGTTCCTTCAAGCTTACGAAAACGGAGACCGTAGGGGAACGAGTTCGTAATGAA

*5’-UTR region has been highlighted in yellow colour.

*MtLAP1pro*

CAGATGACCATCTATGTAAGTCCGAAAAGAGCATTCCAAAAAAGTGTACAGCATCATTAAACACAGTCCAAATATCTGCTCCCCCCGTGCAATACAAAAAGACCCTATCACTTCCCATAGGAGTCACAATAACATTCGAAAAACCTGCATCCTCCACCCTTTGTTGAAGAGATAAAGATGAGTCTC**CCGAC**ACTACGGTAGCCACCATACCTTCACTAGCCCAAACACGGTCCTCCGGTTTTGAATTATACAAAAGCGGACAGAAAATATCAACTTTCTCCTCCACCTTTGAAATCTTTTGCAATCTGTCTTTCTTCCCTTCAACCTTTTTCATCTCTTCCTCCACCTTCAACACATCTTTTAAAACCACCTCTCCCTCAGCTTCAACATTCTTTTCTTTTTTAAAATATAATTTTTTTTCTCCTCTTCTCTACCGGAACATCCACCAAACCAACCTTCACTATGTTTTTTCCAGCCTCATCACTGACCTCCTCCTCCTTCTTACCCAACCGCACATTCTCTATACCAACACCATGGGTTACCACCACCGGCTGAAACTGTCCAGCCTTTCGAACAACAATATTAGAAGCCTTAACCTCCTCATCATATTTAGCAAATCTATCAAACCTAGCTTCCCTTGCCCATACACGATAATCATCAATCCAGATATTATTCAAAGCATGTTCTAGCTTGTCTCTATTCTTTA**CCGAC**TCAAATCTAACAAAACCATAGACTTGACCACGCAAATTCAGATGTCGAGCAATGTAAATATCGGAAAGAATACCGCATACCTCAAAATATTGTCGCACGCGGAACACAGGCAAAGACTCTGGAATATTAGTAAAATAGAAAGTTGCCTTGCTGCCAAAATCTGTGTCACGAGAGTGTTGCTGGAAAACACCTTGCTGCACCTGCGCATTACGCCTAACAGGTTGCTGATCACGTTTTGCCCTAAATGCACCAGATGACCGCTGTTGAACCTGCCTATCCCTCACCTGATGTCGCGCTGATCCTTGTCCCTTCCTGTAAGATTGTCGATCAAAACCATCTGCATAATCCGAACCTCCTCGTCGCTCTTAAATCCTAGACCTGTCCCATCCTCGATCCTCCGGTTTATTTTTCAGTTCTATTCGTTTCTTTTTTTTACGAATGAACTACTTGTTTTAGGGTAAAAAAATTCATATCTAGTCGAGCGCACCCATTTATCTTTGGGAAATGTTAACGAATATTCGAAGTAGCTATTTACACAAAAATACATGTAATCAATGATTTTGAAATCTAAATATTTTAATTTTTTTAAGAAAAATTGTTTATTTTAGAATGTTTAAAGAATCTCTCGGACACTCATTAACAAAACTCTTTATCTTTTACGAAAAAAGAAATTCCATGCGAGAATATGTGATTTCAACAAACGTGTGTCTCTCATAAATTATTTAATGGTATATCTTTACCATTTTTAATAAAAACTAAATTTTTCTCTAAAAAAAAACTAAATTTATCTATTCTTACTAATATATAAAAAAAACTACCCCTTTCAGCATTTTTAGGTTGGATTTTTCTTTCCAAAAATGCCCTCAATTTTCAATACAAATAACACATGTAACAAATAGATAAAAATAAATTTAAATATAAAAAATATAACAAACACGATATATATATATATCTTTTTTTTTTTGTCAAAAAAAAAGTATGTCTGTTTCTTTTTCTTTATCATTTAAAAAGTGTAACAAACTAAATGAGTATATTTATACTTAAATTTTAATTATCTCAATTATACCCTTAAATAAATTTTTCTATTATAAAGATTGTTGTTCGTGTTATTGAAATAAATTGTCATGTTTAATTTGGTTTGTTATAACTAGAAAACAAAAAACATTTATATTTTTAGTTTTAATCAAAATCAAAATTTCATAAAAAAAATATAGTTCATAATTTTCAAATTTTAGCGCAGTAAAAAGAGTGGAAAGATGAGCACGTGAGTGTCTGTCTTTTTGCTAGTAATAATAATAATAATAAATATAGAATAAAAATAATATTTTATTAAAATTTTCTAAACAATAGGAGGATCCTCAGTTTTGTATTTTCTAATTTTTGTTATCTTTGTGTGGTGTTGCTTTGTTTGTTTGATTCAGATTAATTATGAGATCAACTTCGGTTAAGTGTAGATTCAATCAATGATTTTGGCGTCGTCAATATTGGAGATCTGAGAGTATGGGTTTTGCAGAGATTTATATAGTCATATTTGTAGTCTTTTGTACTTAGTAGTTGAATGCTATTAACATAGATGTTATGAGTTTATTTGTATATCCATCGTTGTTGTTTTTAACGAATTTGTATTGCATATTGAATGAATATCGTTTATTTTTAGTAAAAAAAAATTTGAATTAACTACAATTTTTCCACATGATTAACCGGATGAGTAAATACTAACCGTGTTTGACACCAATCATTTAATTTATATGAGGGCAAAAATTCATGTCTATGGACAATTGCAAAGGAGCATATATCTATAT

*MtDFR1pro* (no CCGAC/GTCGG)

GTTTATTCTCCCGAAAATGGAAAATCTATTCAAACTGACTGAAAGAAATTACAAACTTATACGTATTATACTACTTGTTTAGATTAATATTTACTCTTTTTCGTTCCAGTTTAAATGTAAGACTTTTAACTCTTTATCATTTAGATTAAATCAATGGACCTATCAAAACCTCACATTCTTTTTATATATTTTTTTTTTTTGAATAATGGACTTTTAACTCTTTATCTTTTATTAAATCAATTGACCTATTATCAAATCCTCAGATTCTTTCTTAGATCTTATGTTTAATATCTTAAACTTTTTTTAGGGATGTTTAATATCTTTAACTTGAGAGGTGACAAATGCATTTTTTGTTGGTGAATTTGGTTTTTTTTAGGGGTTTTCGTTTTTCTATTTTAAATTTTATTATCATAAAACAACTTGCATGTGCATTGTCACCCACTAATGTAACATGGCAGCTCATTCTTTTATTCAACTTGACAAGTCACCTAAAAATACCTCCAGTTCAATGGTGGATTGTATTTACATTTTCAAAATTTTGAAAATTGAGATTATATTTTCAAACTTAGAAACAACCAAATAGTGAATAAAAAAAAATAAGGGACCAAAATAAATTTTAAGTCAGTCAAATTAAATGTAAATAAAATATTCAACCTAATGTCCCTAAAAAAAAATAATTCAACCTAATGTTTTAACTTTTAACCATGTTTCGAACCAAAAAATGTATTAACAAAAATGCATCAAAGCTTACTAAATTTATCAAAAATAAAATAAAAAAAAGCTTTTAATTGAAAGTTTATTTTTTATTTCATTTAATTTATGATTCACAATATATATTTTTGAATCATGATTCACAAAATAACGTAACTAAGAGCACTCACATCCATGTCACCCATATGAGTAGTATAAATGGGTCTCACATAATATTTTATATTACTTTACATTTTCCAATTATTTAAACCACCCAACTACATTTTTCTAAATATCCACACGACTCTCCAAAACTCTAAAATAGGTCTCACTAATCTTACAATATATTCAATTTTATATTTTAAACTATTGATAAAAGGTGAAAAAGGAAAAGAATGGCACCATTGCCATTCTTGGCACCGTAGTGTCCAAGCAGTATCCAAACTGACACACACAGGCACACGTTGGCAATACCACTGCTGATATTTGTGCTCTAAGCTATCATAATTGAGGATTCGAAAGAAAGAAAAAAAAATTAGGAAAGAATCAATTTCATTAATCAAAAAATCATATATCATGATTTTTTTAAGGGAAAAAATCATTTTTTTTTAAGGAAAAAGGGAAAAAATCATATATCATGATATAAATATTCAACCTAATTACCATATTAAAATAAGGCAGATACACATATACTATTCAACATGTAACCAAAAAACATATACTACTCAACTTAAGCGATTCTCCCTTTTCTTCACACCACACACCCACCAAAAAAAAAGAGCAATGCTATATTTCTAAAAAAATAACGTCAAGAATTCTTCAAGAATAAAGAACAGTCAATCACAATATATTTCAATATAGATATTTGATGTGATCTGACATAAAAATAAAAAAATAGATAAATTATGAGAGTAACTCATGTAAAACTTTATTACAAAGATACGTTTTTTTTAATCTATAAACAATTTTGGTGTAAAATCAATCTAAAACGCAATTTTTTTCATCCTATTTTCTTATTTTTTGGGTAACATTTGTTGATTGGCAGGTGAAAGTCATCCTCTCCCTCTCATAAATTTGTAGGCCACGCCTATCCCTATTACTTTTTGTTTGGAATATAGCACAAAATCTTCTATAAAATCAGACCCACAATTCACACATTGTCTCCATTCCACTTTCAAGCCAA

*MtDFR2pro* (no CCGAC/GTCGG)

AGAAACTTCCAAATTTTATATATACTAACAAGATCGGTTCTCAATCAGTAAAGGTCAATTTATTAAGAAAAATGGGATGATACATAGAAAATATTGTTTTTGTCACGGTGAAAAAAGTGGAACATTAATTGTACCTTATCCTATGCAATGTTTGCATAGATTAAATTCAATTCAACCAATGGGTTTAGATTGTAACTTATTTAAACTTTTATTTATTTGAAAAATATAATATAATATAATACCACCATCCCATGCATCTGGCCTTTTTGGCAATATATAAATTAAACATTGAATCCTCAATTATTCTAGAAATTATATAAATAGGCGAAGGAGTTACGTGATGTGTTGTCACTGATATTGTTGAACATGTGGATAAAAGGGGGACAACACCCTCATCCTAATTGTGTTGTATAGAGTTAGAGGACCGGCCATTTACTTGGATTAAAAGTTGTGGTTTTGTTAACATTTCAATATTTTAGAGATGAAGTTGAAGACGAATCATTCGGATTTTACTTACTTACTTACTAGTCTAACTTTAAATGATACTACCGGAAGGTTGTCATGAGCTCCCACTTACCAGCTCTTGACTTCATTCCTTGCTATGCCGGCAGCTGATCACGTGCTTATTCAAAAAATCTCGTGACTTGTATTAATTGGATTTTGGACTACTATACTTATTTCCTAAAATTATTAATTCATACAAAAATGAATTAGGTCAACTAAGACAAAGATAAAAAAATAAACCATTACGATTAACATTGTCAATTGATAAAAGAGAATAAGGTGGTGTGAGAAAAGAAAAAAAATATTATAACAAAAAAGGAAACTTACCTACCTACATAACTTTTTAGTTTAATAGGTCATTTGTTCCCTTAATTAATTTCAAATTTTTATTTTGATCTCATAAGTAATAAAATTCATGTTTTGGTCCCTCAGGTTTGCCTCCGTCGTCAAATAAATCATTTCCGTCAACTTTAACCTCAAATGCCTATTTTTCTAAAAAAATAAAATAAAAATCTCAAATGTCTATTGTGGACCAACATTATAGATAGTCCACCATAGACCTTAATAATTTTTAAAATTTTAACCATTAGATCTTTTCTCATAAAAGAACACCGTATGATCATGCATGTCTATTGTGTTTTATGGTGAAGCAGCAACACCTAATATTATTTAATCTTTCTTCATCTTCTTCATTCTCATCTTCATCATCTTCTTCATACAACCTCATCATTTTCATCAATCTTCAAACTTAACCCAAAAAATCAAAAATTAATATCACAATTTTTTTTTTTAAAACTTAATCTTAACCTCTTCTCTTCGTTGTTATATGACTTATGACAACTCAAATGTTTATAGCTAGTAAATTTAGAAAATAGAATTATCGTTTTTGTGAGTTTAACTCAGCTGATATAGATATTGTATAATATATGTAGTGTTGGATTTCAAACCGTGAATACTACACTTATTAATATTAAAAATGTGAATATTGAAATTACAACCCTAAGTAAATATTATTTAACAAAATAAAAGAAATACCTTAGTGGGATCACAACCCTAAGTAAATATTATATAGACATATAAAAAATATATATACATCATTTACTTATTTTAAAAATACGTATATTTAATTCTTCTTTGCGTGTGTTATAGATGTGCTTAAATATTGGGATCACACTATACAGACCTCCGCACCAAGGAAGATCCTTCACGTCACTTCATACATTCCTTAATATCCTTACATTTCCTTCACGTCACTTCATACTATATAAACCCCTTACATTTTCTTACTCTCATCAGCTCAGGTTTCTCTTCATTCCTCTCTCTTTACAACAGCTTCTTTCTAAATAAATAGAGTCACCCACCACATACATTGTCTTCC

*MtANSpro* (no CCGAC/GTCGG)

AACTCTCTGTTCATTGCTTTTCATATGATTGTTTGCATGACTATTAACCCTAACTTCTTGCAATCACATAAATTGATCTTAGGAATTGAATGACTACTAACCTTAGCTTTTAATCCTGAAATAATAATTCTGACATCGCTTCCGCGGTTATTCAATCGGTTTAAGAGATTAAAAGGTTATAACTTAGGAAAGGGTTTTGAGTCAAGAGATTGATCAAAACTACTTAGTTAATTTGTCGTAAAGAATGAATAGTAATTATTAATAAAGGGAGGTAAACTACTCATATAAAGCAATAGCGCATTCATCGGCTTAATCATCCTTTTCAAATCATCTTTCAACAACTTTTATTACTTTTTATGCACACAAACCTACTGCCTCAGGCACTTATAAAATTTACACATCCTAGATATTAACTTTATTGCTAAGTTGAGATTAACACAGTCCTCGTGGATACGAACTTATTTTATTACTTCGATAGTTTTAGTACACTTGCTAAAATATTTATCAATGATTATTGTTGTGTGTACATAAGACCCCACATCTCACCTTATTCTCGTTGTTTCACGAAGTTTTTGGATACTCTTTCTACACAATAACTCTCTTAGAACATGCTTCCTAGACTTTAATTATGTTGGTTATCGTGAACCATAAAACAAGGAGGTGAATGTCCTTGAGACTTAATTATCTGTTCCACACGACTTTGATTAATTTTGACATTTTCACGCTCCTTTGTGAAATCAAGATATATTCAAATAATCTATCATGGGATGTTAGTTCAACTGATTACTCGCTCGACTACCCAACATCGGCTAACCCGTAATTCGATTATAGACCTTTTTCAGCTAACAAATTAGATTATTTTTATGTGTAAACAAATACAATCCATAAAATTGTGACCCCACTCCCTTCTCATTTAATAAGGAAAAATTTCAAGTTAATTTAAACTTTATATAATCATCCTTCGATATTAATCATAAACTTGATCCCATGAGGAGTCATAAGATTTATCAACATTTAGGATTAAGAACTTCCGGATAGGGATGACAATGCGGGTTAGCCCACCTCGTTTAGGCTCGACCTACTATTAGCCCGCAAAAACTGGGGTGAGGTGGATCAACTTGAGATGGTGGGCTCGCACACACCGTTGGGTTAGGGGGATGACCATCTAACATTTTTTTTTCTTTCGAAATTCTTTTTTTTTCTTATTGACAAATGTTATTAGAAAATATGCTACTGTTGTAATAAAATCATTTTTATTTTCTCATGTTCATTCTAATCCCGTCTAAAGAGAAAAACAATTGTCTAATAAAGAAGTGTTCAATTATTTTTAGGGAAAAAAACTTATAAATTGAGTCACTTACTATGATAAAAAGTAAATAAATACTTTTTGATCAGGCGGGCCATCCTGCCGCATTTGGCCACGATCCACTTTTTTAACGGGCTAAATAAAACGGACCAACTATAGGTGATGAACTCAAAACCTTAGTCTAACCCACCTAAAATAGGAGGTTAAACAGACTTGTGTGATAGATAAGACTCATTCTGCCACCCCTACTTTCGGATATACTAGATTTTGATTTGATTTCCTTCTTTACACGTTGTTTGAGTGGGAAAGAAACTAATCTAGTTTGATAGGAATTTTGGAGGTTGGGGAAAGACGTAGTAGTGCACGTAGCAAAACTAAAAAGTTAATTGCGACCGTTAAGTTAAGGAGAGTAGCACGAGATTTGTTTTAGCTATAGCTACCGTTCAATCCACGTGCATAGATAAATCATGTTAAGTTGAGTGTTCGCTCATCACGTGTCGCATTATTGGACTCATATTATTTACTTAAAAAATATATATATTAAAAATCATACCATATGTTTATGAATGGGATATTGATCTGTCCTTCCTAGTTGAAAGTGAAGTAAGTGAGGTTGGGCAGTCTATAATAACTAATTGTAAATTGATCAATCAAACATATATAAATAGAAGAGAGAAAGAGAGAAAATATATTATATCCATTATAATAAAAATTATACAAAC

*MtANRpro* (no CCGAC/GTCGG)

AATATGGCACAAAGTTTAAAATATTTTATTTAGTCAATATGTCGCAAAGTCTAAACATAAATTCTATTTTATTTTATTCTAATTTATTTATGCACTTAAAATAAATTCGATTTTATACTAATTCAATTTTCATTCTAATTTATTTATTCATTAAATACAAAATTAGAAAATAAAATCGATTTTATTTATTCAATACATAAGCATAAACATAAAATAAATGTGAAAATAAATGCAAATAAAAAACAAAATTAATAAAAATAACTCTCATTTTATTAATATGATTCAAATATTACATTATATAATACATATATTTTACCCTCTTCGGTATATGACATAGCTCTTGGAGAAAATAGTGATCCGTCCTGTTGAAACATCTCTAAATTCGTGTCTGTGTATCATGTCATGTTGAAACATCTCTAAGTTCGTGTTTGGATTTACGGTGGAATTGTCAAAATTACGGTGAGCTCACCTGATTTTGACAAATCAACCCCTTATATCCCAAACATACATCACAATTATATAATATAAAACTCTCCTCGTTTGGAAGAACTAAAAAATTGTATTTGAGTAGGGTTTAGGGAGGGTTTATATGAATTCTTTAAATATAATCTTTGTGTTTTTGAGTAATAAATATAATCTTTGTTGTTATAGTATTCTTAAAATTAAAATATATTAATCATAAGAATTAGTTTACCACCCTCTAAAAAAAATACTCTTTTAAAAGAAGTGAATGATTTTTCAATTTTCCCTTATATTTCTCACCTCCAAAGCCCTTCCTTCACTTTTCTTCAAAACTCACAAATAAAGTTTAAGACGGTCTTTTTTCTCTCCGTTGATACACCTTTCTTTTCCTATTTATGTTTTTTTTTTACACATTCCTATTTATGTTATGTTATACTATCATACAATACAATTGAAACATTTTGAGTGGTAGATGGATGCATCCGAGTAAGGTAGACTAGGAGGAGTTTTTCCAAAAAAATAATTGAAATGTTTATGAAGAAAAAGCTATTAGGATAGAGGGGATGGTGAAACTTTACCAACTTTTCAAGATACTCATTGATGTCATCATGGATTATGGTGTAAGTTGCTAAATGGTCTGCTCATGAATTTAATAATGGAACTCTATAACGAATTATGAAGCGTTAGTTAACAAGACATAAAATACTTTGTAATCAACTTTTTAAAACTTTAAAAATTGTGTTTTTCATTACAAAACTTTTATTTTTTACTTCCTAGCATGGTATTATTTTGGTCCTTGTATTTTTCTATTTTGGTAGTCTCACACCAATATTTCTCCAAATTTATGTTGGATGTATCATTCCATAGCCAATATAATTTTTCTTTTAAATTATCTTTATTATAAAATGTAGGTAATAAATAAAATTATGCAAAAAATATATTAATGAGTGTCCAAATTTGTAACTGTATTTTAAAATTGCTATAAAATATATATTTTTTTTAGGCGCAAGATACTCTATTCTCATTATAGAGACTCTCCTTTTATCATAAAATTACTGAACTCTATAATACATTGGTACGTAGGCTCGTAGATTTGATAGGTGGTGAATTTCATTCAATGAGAGAATTAATGTTAATGAAAAATTAGTATGTATAATAGTAAAAAATGGAAAGTGAGGTTTTTCAAACTTCTCAAAGACTTGTTGATATTGAGTCACCGGTAGACAACAAACTCCATCACGTGCTCACCTTAAACACCAAAATGTAACAGTTTGATTTGTTCATGTAACTGAACAATTTAGTGTTATTTAAGAACTAAATAACTCTTCTTTCATGACCATAAAAACTGTTTTAGTGTGTTTAAGCTTCATAGTGAAAGAGAGTGTGTAAATTAACATTTGATGGCTAGTATCAAACAAATAGAAATAGAAAAGAAGAAGGCATGTGTGATAGGTGGCACTGGTTTTGTGGCATCATTGCTGATCAAGCAGTTGCTTGAAAAGGGTTATGCTGTTAATACTACTGTTAGAGACCTAGGTCTCTCTCTCTCTCTCTCTCTCTCTCTCTTAACATCTCATTGTATAATAATAACTGTTATGAAGCACATATACTCCTTGAATTAGGCATGTCCAATGTCATTTTCTCAAGTTATTAACGGTGTCGACGTGTTCGTGTCTGTGTTATATGCGGTGTCCGTGTATGAGTCTATGCTTCATAGCTGTCTATATATTCATGTTCATGTTTTATTATCTAAACAGTGAA

*MtLARpro* (no CCGAC/GTCGG)

ATGAAGATGCAGAATCCCACTAGATCTTTCCTGCACCGGTCCCAAGGAAAATCTGGGAAAGTAACTTTCTGATGTATGTGGAAAAGCTCATACTTTGGTTATTGCCCAGAAACTCATATGAAAAGCATATGACACGCCCAGAAATTCATGTGTTTATTCATACATGATGAACCCAGATGAAGATCATCATGAACACAACAACATTCTGATGTTCATCAGTGATCAGAATGTTTGCTCATAATTTCAAGTAAAAGCTTGTGGGACCCAGGACACGTGGCATAACACCATTGGACACCCTACAACGGCTATATGAGCCCAAAGACTCTATAAATAGAGATCATTGCCTTAGCAAAACTTGCACCATTGTAAAGCATACAAGAAAACAATATGACTTTGTTTGAAGCTCTTGCAATTGAAATTAATCAATCTCTAAGGCTTTTGTTTCCTTAGTGCACATCAATACTCCTTGTTATCTCTCTCTCTCTCTCTCTCTCTCTCTCTCTCTCTCTCTCTCTCTCTCTTAAAGATAACTTCTTCTTCCTCCTCTGTTTGATTCACAAACATTCAACCTCCATACAAATTGCAACAAAGTCTCATCTAGCTTTAGGGGTACTAAAGTGTTAGTTTGAGCGAAGGTTGCAAAAGTCTAAGTGGTTAACTTAGCAAGAAGGTGCAAGCCTGCTGAGGCTTAGAGAATACAGAATTGTAATTGTTCTGGTGAACAAAGATTGTAAGGTGCAGGACTTGAAAGGTTATCTCAAGCATCATAGTGGAAACTCTCACAAGATTGTGAGGAGTGGACTAGCCCACGTTGGGTGAACTACTATAATTCTCTGTGTGTTCTTTCCTCTTTCTTTATACTCTTTAATTACAGTATACACAACACACACTTACACATTTACTTTCTTAAACAGTTTCTTTGTTTATGAACTTCAGAACGTTGTGAAGTCATAAAGTTAACTTAACTATTCCAAGTAAACAACTTGAGAATCACTTTTAAGTTTCAGGATAATTTTTAAAGGGTCACAATTCAAATCCCTCTTCCTTATGACTATTTTATCTACTTCAAGTATTTCTATCCAAATAGAAATGTGAACACTTGCTAACTTAGACTTTAATGTTTTTAGACCAACCTAAAAATATTAAACTCTCAATTTAAAATAAAGGACAATTTACACACTTTAGTAAATTGCACTCAACTTTTATTTTTCAATAAAACATTCCAAAAATTGTTTTCAAATTTACAAAAATAATCTTCTGTTTAAACTTTTGTTTAAACAGTAATAAGAGGATATTTTTATCCAAATAAAACTTAATATGACAGCTTATAACACTCGATGCCTTGCTTTTTTTCTTTTTCTTTTTCTTTTGAGGGTTTTGGTTTAGTCTCTCCTCTTTTACGTAATTTTTTTTTTTTAATAAAATTTCGGCGTGAGTAGTAGAGGATATAAGAGTTTCTTTGAGGTGTCAATTATATTGGAACATCTAAGTTTCACCTTAATGTTTCAACATTCTCTTGGTTGATATATATATATATTCTTTTTATAAAACTGAAATTCGCCGGTCTTTTAAAATTCTGTAGTTTTGTTTTACCAAAAAAAAAATTGAGTAGTTTGGTCACCTAATCTTTTTCTTGTTTTTTTGACACATGGGACCTAAACTTCAAATCATCAAATCCAAACGTTCTTGTGCATAAGGGTAGTCAGCAGTAAAAAAAAACGCTACATGTGTTGAAGAGTAGATGAAAAACACCCTACCAATATTCCATTTATGGTCATAAATATCAAATAATGTAACGGATACATTGATTGTTCTATATAAACAAACTAGCACCCTTCGATTTTGCCATCTCATACCCAAATTATCACAATTAACTCTTCCTCCAAAAGT

*MtCHS2pro*

GTTATTTTATGATGTCTTGTAATTATGTCAAGTGTAGGGCTCTAAGTAAAGGAAGTTATTTCTACTTTTAATTTAGACCGTAGGATTTATTGGTAGTTTTTTCTGCAAAATCATTGATTTCTGAAAACTCCGCAACACCCACGCCATGGGAGAAAACGGGTTCAGGAGTGGACAGATAAACGGTTTCCAACCACACCATTATATTCCCTGCAATTTAATTCCTTTATTACCGCTTTATTTTCCTTTTGTTTACTACATTTTGCATTTTTACTTTCATTTTCTTTCTTTTGAATGCTCATTTGAGTGTTATTTCTTTTGAGTCGTCATTTGATAAATTGTTTTATTAAAGTTGGAAGCAATTATAACACTAATGAGAACGATTGGATTTGTTCTCAACAAAGACATAAGGAAAGAGACCATAAATCGCTAAACTCAAAGAACCTCAAAAAGGACTTTAATTCGGTTCTTAACAGTTTTGGATAATATCATTTGTATTGTCAAAAATCAGTGTAAACAGTTGGACAAACATTATTTGGAGATGGAGCTCATGCGCTCATTGTTGGTTCTAACCGAATACAAGAAATTGAAAAAGTTATATTTTTTATGGCATGGACTATATAAACAATTGCTCCACGCAATGAAGGTGGCCTTCACTTCGTGAAGTTGGGCTAACATTCATGTTCGTGGGATTTTCTCAATGAATATTGATAAAGCATTAACTGAAGCATTCCAACTAGGGGTGTTCGCGGTCCGGTTTGGATCAGTTTTGAGGTAAAATGTCAACCGATCCGAAATAAATATCACATGTGGTTCGGTTCGGTTTGGATGAGTATTCTAAAAAAATCCAAACCGAACCGATCCAAATATATGCGGTTTAATTTGGATCGGTTTTTGGACATCCAAAATGCAAAATATATTTTTTTAGTTACTCTTTTTTGAAGAATATAATTTTTTTTATTAATATTAATATTACACAAACACGGTGAAATAATAGGTTTGATGCAAAATATATTAAGAAGTTAAAATTATTGTATGAAAATTATCAATTATTTTTTAAACATATGTAGTCATAATAATATAGATTAAATTAGTGCAATATATATGATTAAAATTTAACAAGCAACATAATAATGCAATATATAATACTAATAAAAGAATATATAAGTATTAATATTTATTTTTTCGGTTCGGTTCTGTTTAATTCGGTTTTGAAAAGGTGATCCGAAACCCGATCCGATCTTAGCGGTTTTTACAAAACAATATCCAAACACATCCAAATATATTCGGTTTTTTACAATTTTCGATTTTTTTGGATCGATTTGCGGTTTTATTTTGGATTTGAACCCCCCTAATTCCAACCATTAAACATCTTCAATTACAACCCAATCTTCTGGATTGCTCATCCGAGTGGACCTGCAATTATAGATCAAGTTGAGCACAAGTTAGGCTTAAAATATGAAAAATGAATATCACTGTGCCGTAAACCAAAAGGTTTGAGAGTAAAAATGGTCCTCAAAGTTATCATGAGGTTATTTGACATATCTGCCACCCAACTCCAACATGTCACGGACACAACCAGAAAGAAGCCATAGTATTTGTTGAGTTACACTTTCAAAAACACTACACCACATTTCTAGGTATAGTCGGTTTTAGATGGCTTATTTGAGGTTGGAAGTTGGAACTAAAAGTTTAGGAATTCACTAGGGAGACAACACACGAACTCTTAACATCGATTAGAGACAACTCTTTTGGCACATTTAAAGTCTCATATTATTTGAGAATCAAGGCAAAAGGATGGTCTATTGTCTATATACAATATCCACACTCAACCCAACGAAATATCCATCAAAGTCCAATATATTGAAAGCTTTCCATAAATGTCAAAAGTGTTTTAGTCAAACAACGTCGATGTTAAGCTACCGTCCAATTAATATCAACCATGTCCCTTTCCACGATCGAGTTCAAATTGCACTACTATAAATACACATACAAACTACTACGTACCACCATATTATTCTATGTATGATCTTAAAAAGCACATTATTATATTACTTTTTTGCTTTATAATTAGAAAACATATATTAAG
